# Somatic selection shapes mutation accumulation in *Citrus sinensis*

**DOI:** 10.1101/2025.09.24.678405

**Authors:** Taylor R. Beaulieu, Emmanuel Ávila de Dios, Danelle K. Seymour

## Abstract

An organism becomes genetically mosaic through the accumulation of somatic mutations. Genetic mosaicism is a commonality of multicellular life and has been studied extensively in humans due to its associations with aging and diseases. In humans, somatic selection shapes the accumulation of somatic mutations, with strong signatures of positive somatic selection in cancer cell lineages. So far, evidence for somatic selection in plants has been inconsistent. The evolutionary implications of genetic mosaicism in humans and other animals are limited by early specification of germline cells, preventing transmission of somatic mutations to progeny. In contrast, many plant lineages reproduce asexually with clonal progeny derived from vegetative tissues. We describe the patterns and processes shaping somatic mutation accumulation within a single, 149-year-old historic sweet orange (*Citrus sinensis*) tree and within a clonal lineage of sweet orange. More than 12,000 somatic mutations were identified in the historic tree and 28,000 somatic mutations were identified across 199 clonally related sweet orange accessions. Both the spatial and genomic distributions of somatic mutations are non-random. The spatial patterns of somatic mutations across the historic tree depend on tree growth and development and their accumulation across the tree canopy recapitulates branching topology. Analysis of the genomic distribution of somatic mutations revealed that the subtelomeres, which are large arrays of ∼180 bp repeats, are mutation hotspots. Finally, there was genomic evidence that somatic selection shapes the accumulation of somatic mutations both within the historic tree and also during clonal propagation.

## Introduction

Multicellular organisms become genetically diverse throughout development due to the accumulation of somatic mutations (1). Somatic mutations arise during cell division from DNA replication-dependent processes, including DNA polymerase errors, and failure of DNA repair mechanisms in response to age-related and environmental sources of DNA damage (2, 3). The result is a genetically mosaic individual with a spectrum of new mutations jointly shaped by DNA replication errors and DNA damage (4–8). Genetic mosaicism has been documented across a range of multicellular organisms, including both plants and animals (9–16).

Somatic mutation accumulation patterns are well-studied in humans in relation to aging, cancer, neurodegenerative disease, and in healthy tissues (6, 17, 18). There is abundant evidence of somatic mechanisms which shape the accumulation of somatic mutations in humans, in which strong signals of positive selection and weaker signals of negative selection have been observed in the evolution of cancer cells as well as in non-tumor cell lineages (19–23). Comparatively little is known about the somatic forces which shape mutation accumulation in plants. Somatic genetic variation due to somatic drift has been observed in clonal populations (24), and models have been developed to characterize somatic drift during branching events (25, 26). The contribution of somatic selection to somatic genetic variation is less clear. Some prior studies find minimal or no evidence for somatic selection acting on genetic mosaicism in plants (10, 11, 15, 16, 27–29), while others identify signatures of purifying selection (24, 30). Despite conflicting evidence from genome-wide surveys of somatic mutations, evaluation of the fitness effects of inherited somatic mutations in progeny from self-pollinations of the same or different flowers on a single plant lends support to a role for somatic selection in shaping mutation accumulation during plant development (31).

Detecting all somatic mutations within a sequenced tissue is a challenge (32), and inconsistent evidence for somatic selection in plants may be due, at least in part, to low numbers of observed somatic mutations in previous studies. Studies of genetic mosaicism in trees are often restricted to the discovery of mutations present at high mutation fractions, a metric that reflects the fraction of cells in a tissue carrying a mutation (10, 14, 16). Previous research has also revealed that high fraction somatic mutations make up only a subset of the mutations in plant tissues (9, 11, 15). The detection of somatic mutations in angiosperms is complicated by the arrangement of meristematic cells which form distinct cell layers, wherein a somatic mutation is expected to be constrained to the cell layer in which it originated (9, 15, 26, 33). When sequencing tissues derived from multiple layers, such as leaves, somatic mutations are then present in only a fraction of sequencing reads. Thus, the identification of somatic mutations requires high sequencing coverage and methods to distinguish true somatic mutations from sequencing errors which are both present at low mutation fractions.

In humans and other animals, the evolutionary implications of genetic mosaicism are relatively limited because germline cells are set aside early in development, minimizing the number of cell divisions and thus the number of somatic mutations transmitted to offspring. In plants, the number of cell divisions leading to germline cells is also reduced (34–36); however, many plants are capable of asexual modes of reproduction in which clonal offspring are derived from vegetative tissues. During asexual reproduction, somatic mutations can be transmitted to clonal offspring and, as a result, genetic mosaicism may have significant evolutionary implications in asexual lineages. Clonal propagation is extensively used in perennial crops to maintain and disperse cultivars, and somatic mutations play an important role in the development of new clonal cultivars (37, 38). During asexual reproduction deleterious somatic mutations are expected to accumulate in a process known as Muller’s ratchet (39). Mechanisms that reduce the accumulation of deleterious mutations include adaptive mutation bias and developmental selection (40–42). Adaptive mutation bias reduces the occurrence of *de novo* mutations with deleterious effects (43), whereas developmental selection favors the proliferation of cells with higher fitness during growth and development, thereby removing deleterious mutations (44, 45).

To explore the patterns and processes that shape somatic mutation accumulation in a single, long-lived individual tree and a descended clonal lineage, we used deep sequencing to discover somatic mutations within a 149 year-old historic sweet orange (*Citrus sinensis*) tree. First, we describe the spatial and genomic patterns of somatic mutation accumulation. The spatial patterns of mutation accumulation in the historic tree are consistent with tree growth and development and are likely subject to somatic drift during branching. There was also a non-random genomic distribution of somatic mutations; subtelomeric regions were hotspots of somatic mutation accumulation. Finally, a sufficient number of somatic mutations were identified in coding sequences to reveal that somatic selection shaped patterns of somatic mutation accumulation within this historic tree. Additional sequencing of 199 clonal sweet orange accessions provided evidence that somatic selection also shapes the accumulation of somatic mutations during clonal propagation.

## Results

### Deep sequencing of 149-year-old historic tree uncovers somatic mutations across mutation fraction spectrum

To discover somatic mutations in a long-lived individual tree, whole-genome Illumina sequencing was produced from 28 leaf samples spanning the tree canopy of the historic 149-year old “Parent Washington Navel Orange Tree” in Riverside, California. This tree is the last surviving individual from the first successful introduction of navel orange to the United States and is the oldest living navel orange tree in the country (46). Two adjacent leaves were collected from seven branches at two locations representing a trajectory from oldest (basal) to newest (apical) branch growth (Figure 1a). Diploid reference genome assemblies improve somatic mutation detection (47, 48) and variants were identified relative to the haplotype-resolved ‘Washington navel’ sweet orange genome (49). Heterozygous mutations in all sampled cells will occur at a mutation fraction (MF) of one when aligning sequence reads and identifying variants within each haplotype of the diploid reference genome. Thus, mutation fraction can be used as a proxy for the fraction of cells with the somatic mutation. All samples were sequenced deeply (mean 41X sequencing depth per haplotype) (Table S1) to allow for the discovery of somatic mutations across the mutation fraction spectrum.

**Figure 1.**
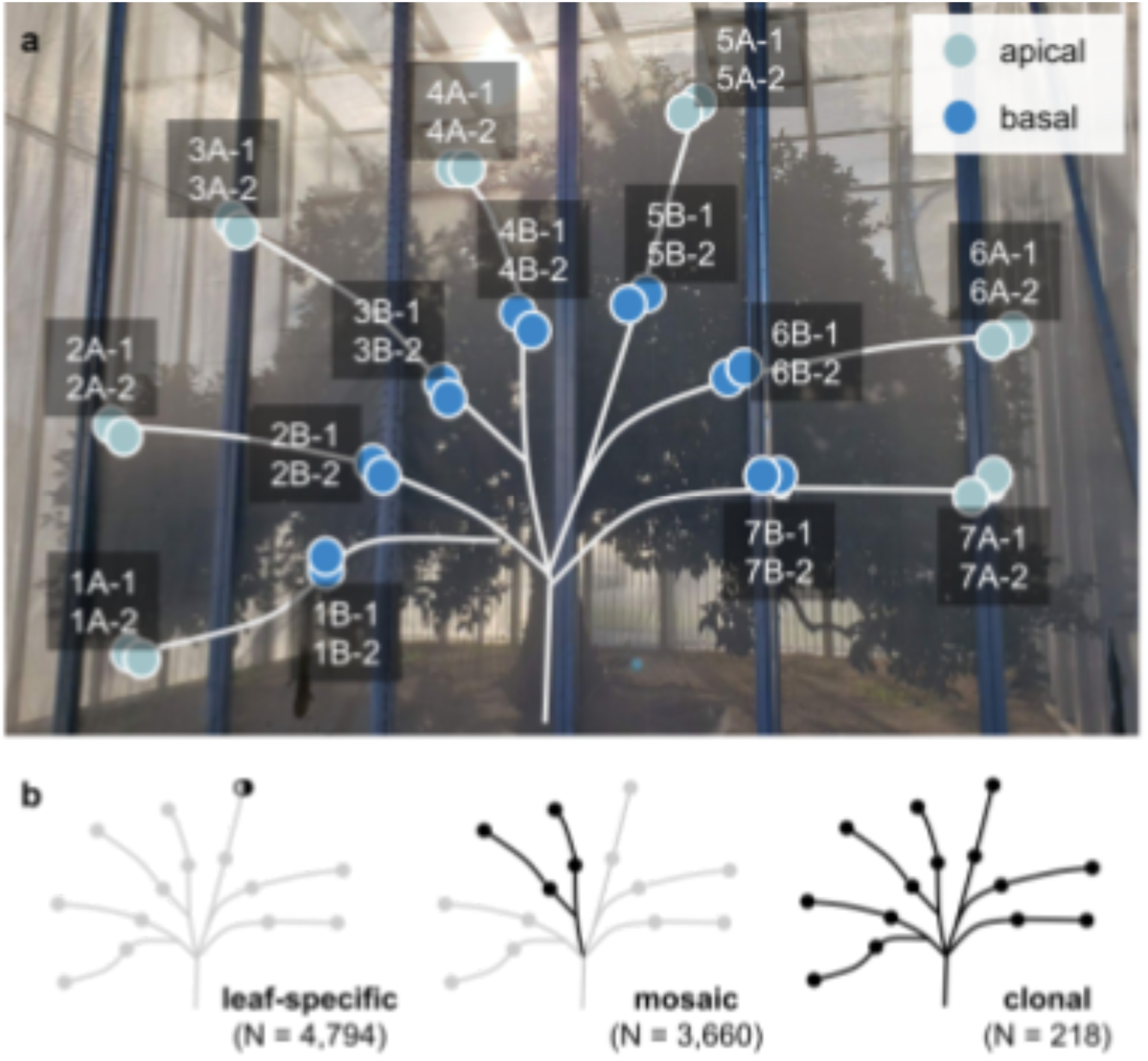
Schematic of the historic “Parent Washington Navel Orange Tree’. **(a)** Canopy schematic of the historic tree Sampling locations at the apices (apical) and bases (basal) or seven major branches are shown. Two adjacent leaves were sampled across 14 canopy locations Branch lengths and sampling locations are illustrative only The historic tree is covered in a protective screen structure for protection from pests and diseases, **(b)** Three sets of somatic mutations were defined based on detection across sampling locations: leaf-specific, mosaic and clonal.

To determine the sensitivity (recall) and specificity of variant detection across the mutation fraction spectrum, *in silico* mutations were introduced at mutation fractions ranging from 0.1 to 1 into 40X (per-haplotype) simulated sequencing reads generated from a diploid chromosome pair (Figure S1). Modification of the expected ploidy (1N, 2N, …, XN) during variant calling has been shown to increase sensitivity for variants at low mutation fractions (18). We found that increasing the expected ploidy from 1N to 4N dramatically improved the sensitivity (recall) of mutation detection with limited sacrifice to specificity (Figure S2; Table S2), and so a ploidy of 4N was selected for variant identification. *In silico* mutations introduced at low mutation fractions (MF = 0.1) are recalled at a lower rate (recall = 69% when ploidy = 4N) than those introduced at higher mutation fractions (MF ≥ 0.2) (recall = 79-96% when ploidy = 4N) (Figure S2, Table S2). The specificity of mutation identification was excellent for variants introduced across the mutation fraction spectrum (MF = 0.1: specificity = 97.5% when ploidy = 4N; MF ≥ 0.2: specificity = 98-100% when ploidy = 4N) (Figure S2, Table S2). There was also good concordance between the introduced and observed mutation fraction for *in silico* somatic mutations (Figure S3), demonstrating that the observed mutation fraction is a reliable measure of the true mutation fraction in sequencing reads.

Based on the *in silico* simulations, optimal variant calling parameters for somatic mutation identification were identified. A total of 12,025 likely genuine somatic mutations were identified in the historic “Parent Washington Navel Orange Tree”. These included 4,794 “leaf-specific” mutations detected in a single leaf and 3,660 “mosaic” mutations detected in adjacent leaves in at least one sampling location (Figure 1b). Leaf-specific mutations arose after the most recent branching event, while mosaic mutations have been transmitted through branching events. A set of 218 “clonal” mutations were detected in all 28 leaf samples, representing clonal variation between the “Parent Washington Navel Orange Tree” and the clonally related ‘Washington navel’ sequenced for assembly of the reference genome (Figure 1b). Together, leaf-specific and mosaic mutations reflect the scale of genetic mosaicism in a single long-lived individual tree, while the clonal mutations represent genetic variation between asexual clones (Figure 1b).

We report a substantially higher total number of somatic mutations than has been previously reported in citrus (73 and 25-32 somatic mutations reported in clementine mandarin (*C. Clementina*) and sweet orange, respectively) (14, 16). Deep sequencing combined with a haplotype-resolved diploid reference genome facilitated robust detection of somatic mutations across the mutation fraction spectrum (Figures S2 and S3) (47, 48), explaining the increased number of somatic mutations detected here compared to previous studies. There was a strong correlation between mutation fraction of mutations detected in adjacent leaves (Pearson’s R=0.93) (Figure S4) and so the mutation fraction of adjacent leaves was averaged for subsequent analyses. The majority (94%) of 12,025 somatic mutations had a mutation fraction of 0.5 or less (median MF = 0.15). Consistent with previous studies in tree species (*Musa basjoo*, *Salix suchowensis*, *Dicorynia guianensis*, and *Sextonia rubra*), we find that the majority of somatic mutations are present at low mutation fractions (11, 15, 30).

### Developmental trajectories shape somatic mutation accumulation across the canopy

The apical samples at the branch tips are separated from the main trunk by a greater number of lateral branching events than the basal samples (Figure 1a). Lateral branches are formed from axillary meristems, so a greater number of meristematic mutations are expected at branch tips relative to the branch base. Indeed, mosaic somatic mutations, which arise in meristematic cells, are enriched at apical locations across all seven branches. Within a branch there are, on average, 1.9X as many mosaic mutations at the apical location versus the basal location (Figure 2a). Leaf-specific mutations are predicted to occur in differentiated cells after the most recent branching event and, unlike mosaic mutations, did not consistently accumulate at branch tips (Figure S5). Based on the transmission of somatic mutations through lateral branching events, the spatial distribution of somatic mutations within a tree canopy should be concordant with the branching topology (33). A phylogenetic tree generated from the mutation fraction of mosaic mutations recapitulates the branching architecture of the tree (Figure 2b), consistent with previous studies of somatic mutations in trees which conclude that the distribution of somatic mutations follows the growth of shoot apical meristems (29, 50).

**Figure 2.**
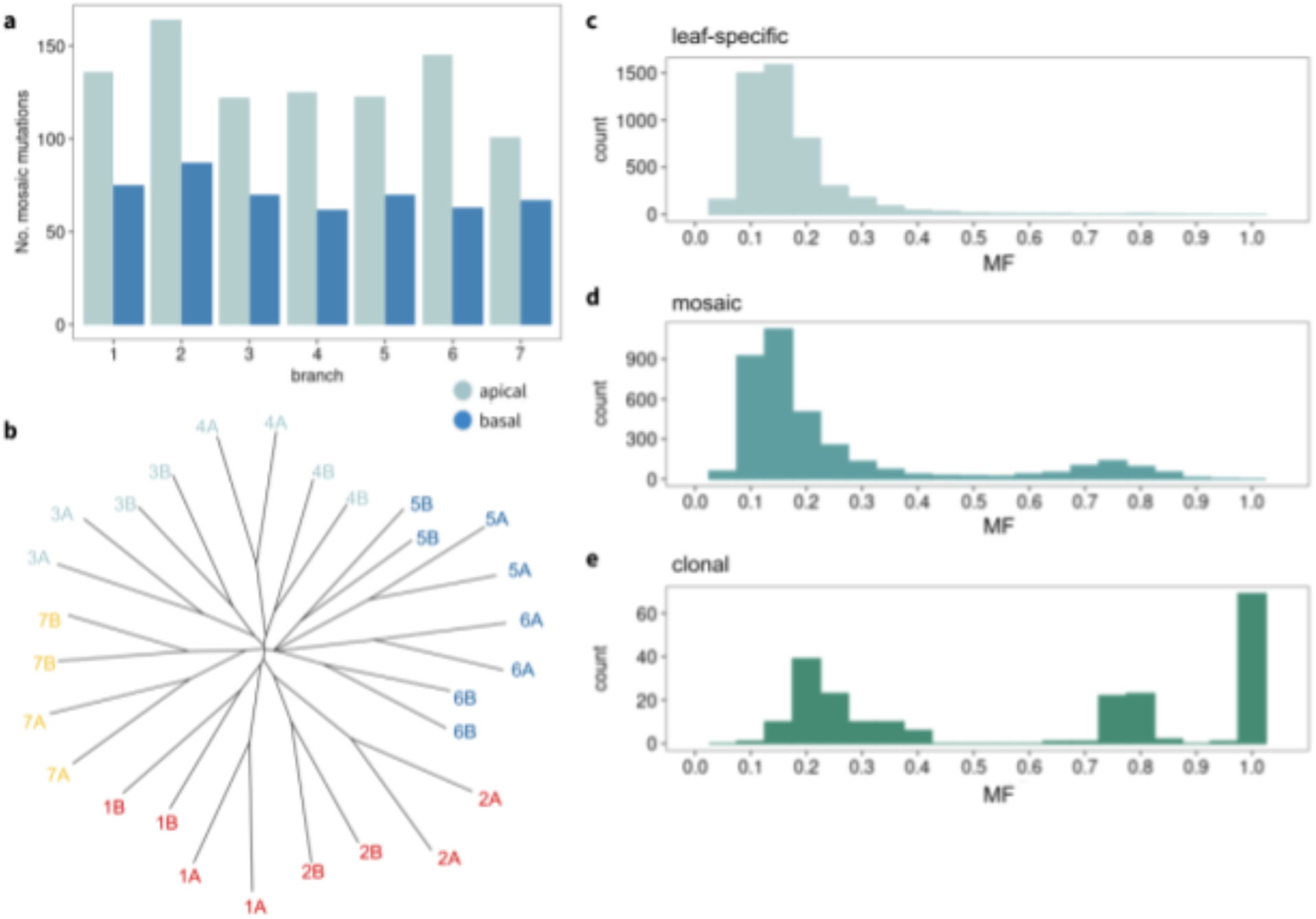
Developmental trajectories shape spatial patterns of somatic mutations. **(a)** The number of completely-detected (i.e. identified in adjacent leaves) mosaic mutations at each sampling location, **(b)** A phylogenetic tree constructed from the MF matrix of mosaic somatic mutations recapitulates the branching architecture of the tree canopy, **(c-e)** The observed MF distribution of **(c)** leaf-specific (n=4,794), **(d)** mosaic (n=3,66O), and **(e)** clonal (n=218) somatic mutations.

Theoretical models of branching predict that somatic mutations can be lost during successive branching events due to somatic drift (25, 26). Low-fraction mutations are prone to somatic drift during branching and, as a result, their spatial distribution may not match branching topology. Supporting this, a study of low-fraction mutations in two tropical tree species found that the spatial distribution of these mutations was not concordant with branching topology (30). In addition to somatic drift, discordant branching topologies could be due to reduced detection of low-fraction mutations, for which recall rates are lower (Figure S2). To confirm whether low-fraction mutations are more likely to have discordant spatial distributions, mosaic somatic mutations were classified based on their spatial patterns as “completely-detected” or “incompletely-detected”. There were 1,237 completely-detected mosaic mutations which were similarly present or absent in adjacent leaves across all 14 sampling locations, and 2,423 incompletely-detected mutations which were present in one leaf and absent in the adjacent leaf at one or more locations. Only the spatial distribution of the completely-detected mosaic mutations was concordant with branching topology (Figure S6). The distribution of mutation fraction differed between the two classes of mosaic mutations (Figure S7). Completely-detected mosaic mutations had higher mutation fractions (median = 0.28) than incompletely-detected mutations (median = 0.13) (Figure S7). Regardless of the patterns of mutation detection across the canopy, the spatial distribution of high-fraction (MF > 0.5) mosaic mutations was concordant with the branching topology, unlike low-fraction (MF < 0.5) mosaic mutations (Figure S8).

Somatic mutations may be chimeric, including periclinal mutations, which are present in specific meristematic stem cell layers, and sectorial mutations, which are present in a sector of one or more cell layers (25, 47). The mutation fraction of chimeric mutations will be less than one for both periclinal and sectorial mutations. The mutation fraction is below one for the vast majority of leaf-specific (99%), mosaic (100%), and clonal mutations (80%). The distribution of mutation fraction for mosaic and clonal mutations are multimodal (Figure 2d-e), indicating either the presence of periclinal chimeric mutations, the effects of somatic drift resulting in sectorial chimerism, or a combination of both. The number of times a somatic mutation is transmitted through branching events influences its mutation fraction (25). The mutation fraction distributions may represent only a snapshot in time, with mutation fraction shifting as the tree continues to develop new branches. However, the distribution of mutation fraction for clonal mutations is also multimodal (Figure 2e). Clonal mutations are maintained through iterations of asexual propagation, suggesting they may include periclinal chimeric mutations. Peaks in the distribution of mutation fraction roughly correspond to the proportion of mesophyll (78%), epidermis (9%), and vasculature (7%) cells in the leaves of another fruit tree species (51).

### Mutation spectrums in meristematic and differentiated cells are distinct

Mosaic somatic mutations must have arisen in meristematic cells, allowing these mutations to spread across the tree canopy. In contrast, leaf-specific somatic mutations occurred after the initial meristematic cell divisions which formed the leaf primordia, and, as a result, are found only within a single leaf at low mutation fractions. Mosaic and leaf-specific somatic mutations were used to investigate patterns of somatic mutation accumulation between meristematic and differentiated cells. Factors promoting cell cycle arrest in meristematic cells in response to DNA damage do not inhibit DNA replication, either during cell division or endoreduplication, in differentiated tissues (52). To determine whether there is evidence of differential responses to DNA damage in meristematic and differentiated cells, we compared the mutation spectra of mosaic and leaf-specific somatic mutations. Leaf-specific mutations are predominantly enriched for C>A mutations (Figure 3a), a signature associated with oxidative DNA damage (4, 6, 53). The elevated contribution of DNA damage to the mutation spectrum of leaf-specific mutations suggests that DNA repair mechanisms are less effective in differentiated cells compared to meristematic cells. Mosaic mutations are predominantly enriched for C>T mutations, particularly in CpG contexts (Figure 3b). This mutation signature, attributed to both 5-methylcytosine deamination and DNA polymerase errors during DNA replication, is commonly observed in both plants and mammals (4, 6, 10, 11, 32, 54, 55). Clonal mutations were similarly enriched for C>T mutations (Figure S9, 3c). Overall, the mutation spectra of somatic mutations in the “Parent Washington Navel Orange Tree” canopy suggests that mutation accumulation in plant tissues is jointly shaped by DNA damage and replication errors, similar to what is observed in human somatic tissues (4, 6, 56).

**Figure 3.**
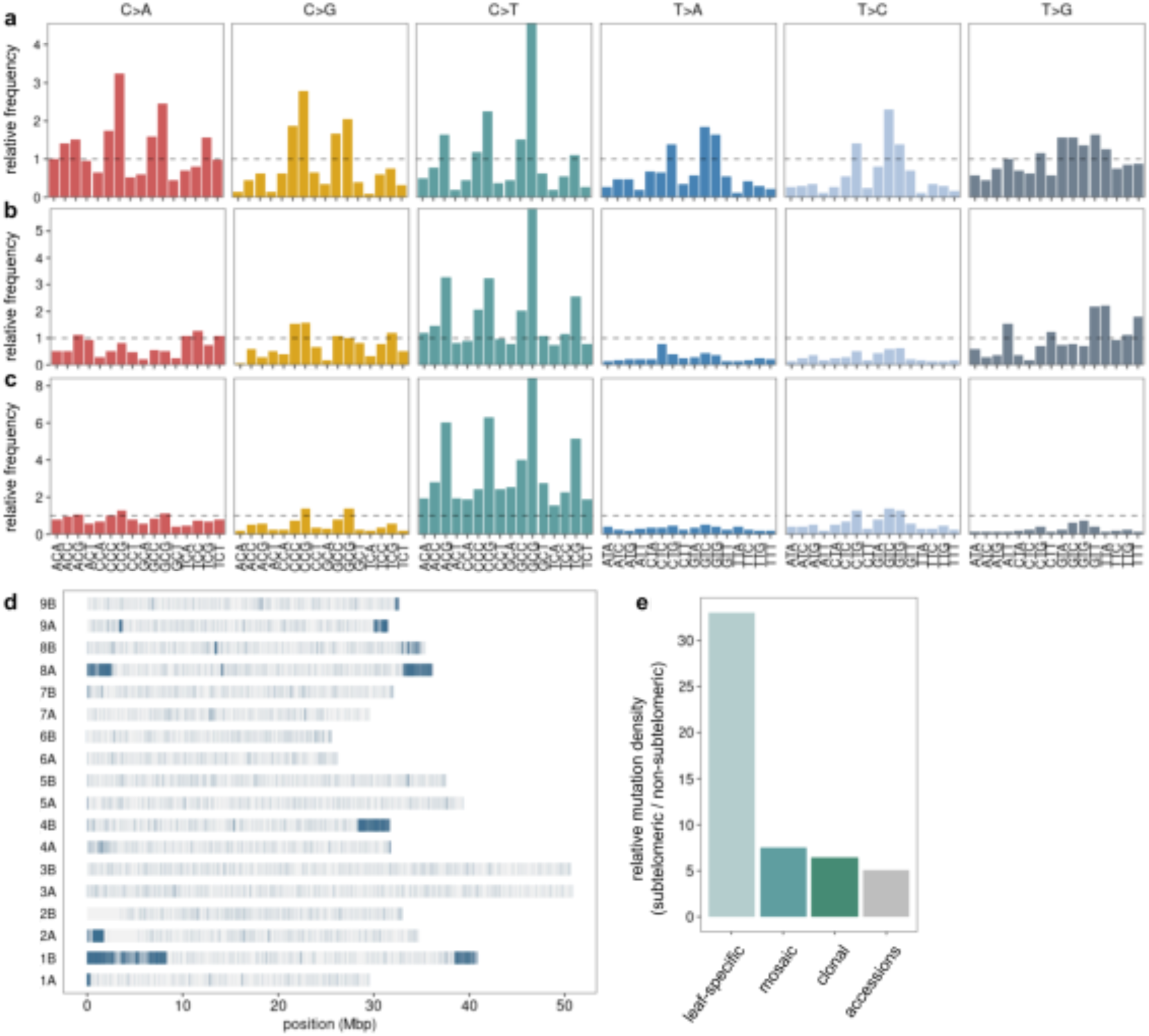
Genomic patterns of somatic mutation accumulation. The proportion of each base change relative to genomic trimer frequency among **(a)** leaf-specific (π = 4.794) and **(b)** mosaic (π = 3,660) somatic SNVs within the historic tree, and among **(c)** clonal SNVs detected in the set of 199 clonally related sweet orange accessions (n = 28,007). **(d)** The chromosomal position of all somatic mutations (n = 12,025) detected in the canopy of the historic tree, **(e)** Enrichment of somatic mutations in the sublelomeric regions Enrichment was calculated as the ratio of mutation density In the subtelomeres relative to mutation density in non-subtelomeric regions for the sets of leaf-specific, mosaic, and clonal mutations detected within the historic tree and the set of mutations delected an six clonally related sweet orange accessions sequenced with PacBio HIFI reads (n = 25,943).

### Subtelomeres are hotspots of somatic mutation accumulation

The genomic distribution of somatic mutations is not uniform (Figure S10a). Instead, somatic mutations are concentrated on the ends of several chromosomes, particularly on chromosomes 1B, 4B, and 8A (Figure 3d) and corresponds to the locations of subtelomeric tandem arrays of 181 bp satellite repeats in the ‘Washington navel’ genome (49). Satellite repeats of 179-183 bp have been previously identified using fluorescence *in situ* hybridization at the terminal ends of chromosomes in multiple *Citrus* species as well as in the citrus relative *Poncirus trifoliata* (57, 58). The highly repetitive nature of the subtelomeres makes their assembly challenging, and as a result, published genome assemblies may not include these regions. Genomes assembled using PacBio HiFi reads, such as the ‘Washington navel’ reference genome used here, may contain these regions. Similar 178 bp satellite repeat arrays were identified in a long-read genome assembly of pistachio (59).

Subtelomeric sequences are hypervariable, with extensive structural variation, repeat copy number variation, and single nucleotide polymorphisms (60–62). We find a higher density of somatic mutations in the subtelomeres for all three classes of somatic mutations (leaf-specific, mosaic, clonal) indicating that subtelomeric accumulation of somatic mutations occurs within individual trees and also between clones (Figure 3e, Table S3). For leaf-specific mutations the subtelomeric mutation density is 33 times higher than non-subtelomeric regions. Mosaic and clonal mutations are also enriched in the subtelomeres, but to a lesser extent than the leaf-specific mutations (8X and 6X higher density, respectively). Excluding the subtelomeres, the number of somatic mutations per chromosome is correlated with chromosome length (Spearman: 0.76; Figure S10).

Because the highly repetitive nature of subtelomeric regions may increase false positive variant calls from poor alignment of short-read sequences, we additionally identified somatic mutations using PacBio HiFi long-read sequencing reads from six clonally related sweet oranges relative to the ‘Washington navel’ reference assembly (Table S4). The six accessions, including ‘Olinda’, ‘Moro’, ‘Shahani Red Navel’, ‘Powell’, ‘Cara Cara’, and ‘Washington navel’ were sequenced to a mean per-haplotype sequencing depth of 40X (Table S4). A total of 25,043 SNVs were identified in one or more of the six individuals across a range of mutation fractions (MF = 0.02 to 1). The density of these SNVs is five times higher in the subtelomeres (Figure 3e, Figure S11), providing further support for the subtelomeres as hotspots of somatic mutation accumulation both within individual trees and also between clones.

### Selection shapes the accumulation of somatic mutations

To search for signatures of somatic selection, we excluded mutations located in subtelomeric regions. Subtelomeric regions are typically characterized by low gene density, few essential genes, and transcriptional silencing (63). These patterns were consistent across the subtelomeres in the ‘Washington navel’ reference genome. Few genes were annotated in the subtelomeres, and annotated genes had reduced length and fewer exons than genes from other genomic regions (Figures S12, S13, S14). There was also an elevated proportion of nonsense mutations in subtelomeric genes among both leaf-specific and mosaic somatic mutations (Figure S15). This is consistent with the subtelomeric gene annotations representing either false gene annotations, or recently inactivated or silenced genes (64) similar to patterns observed in yeast (63).

Somatic mutations can be passed to clonal offspring during asexual reproduction and clonal propagation, and so the signals of somatic selection may also shape the genetic diversity of clonal populations. There were few clonal mutations detected from the canopy of the historic tree (n = 218). To obtain a clearer picture of somatic mutation accumulation patterns during clonal propagation, we examined the genomic distribution and mutation spectra for 28,007 SNVs identified across 199 clonally related sweet orange accessions, including 88 accessions from the Givaudan Citrus Variety Collection at the University of California, Riverside (GCVC) as well as publicly available sequencing data for 111 accessions (14). Similar to the historic tree, clonal somatic mutations were enriched for C>T mutations, particularly at CpG sites (Figure 3c). There was also a 10-fold higher density of SNVs within the subtelomeres in these accessions (Figure S16).

Somatic mutations affecting cellular fitness will influence which cellular genotypes are successfully transmitted during development (45). To search for signatures of somatic selection, we focused on the set of 6,122 non-subtelomeric somatic mutations.

A reduction in the number of mutations in protein-coding genes is expected if somatic mutation accumulation is shaped by mechanisms to reduce deleterious mutations (41, 42). We find that the proportion of genic mutations is significantly reduced for all three sets of leaf-specific, mosaic, and clonal mutations relative to the genomic expectation based on permutations (n=1000) (Figure 4a, Table S5). Clonal mutations identified across 199 accessions had the strongest reduction in the proportion of genic mutations. In contrast, the proportion of genic mutations was only slightly reduced for leaf-specific mutations. The strength of the reduction in the number of genic mutations correlated with the predicted age of somatic mutations, with the oldest clonal mutations most severely affected.

**Figure 4.**
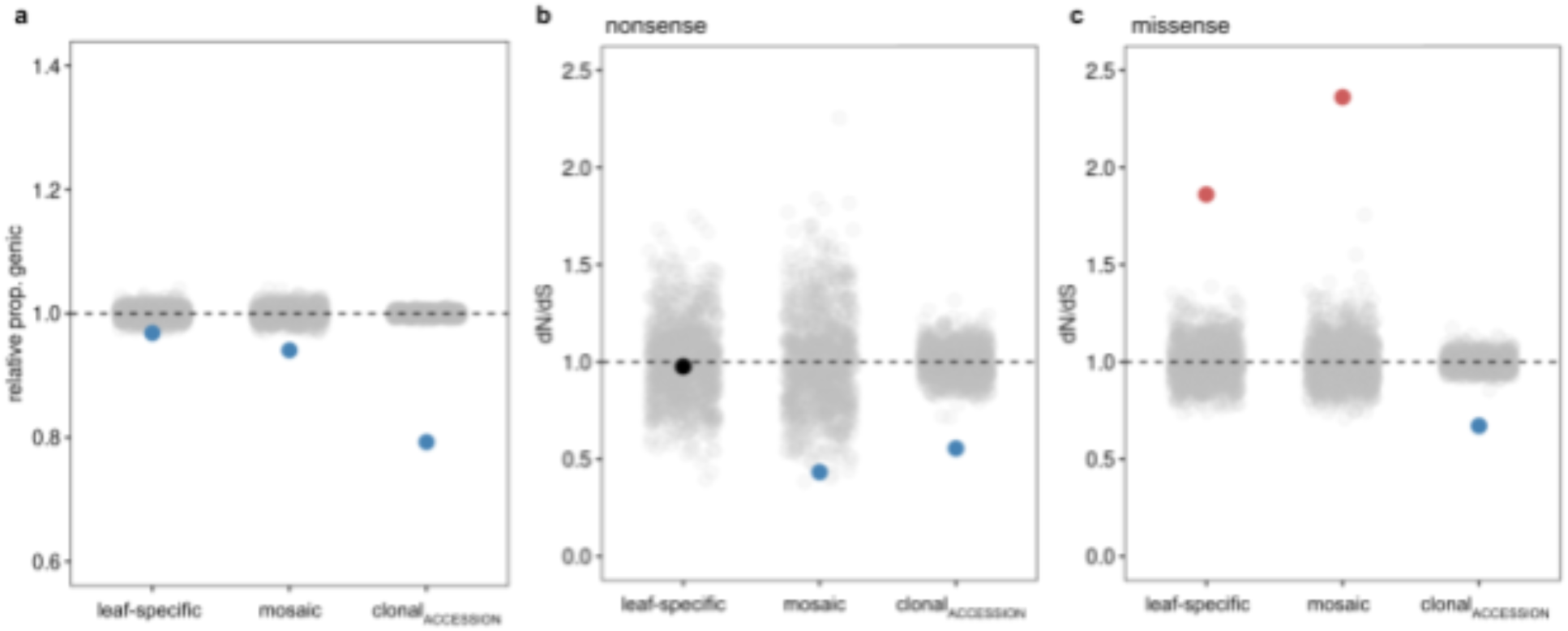
Somatic selection in the historic tree and between clonally related accessions of sweet orange. Leaf-specific and mosaic mutations were identified in the historic tree. For the clonal set of mutations in the historic tree (n=218), no nonsense mutations were identified. A second set of clonal mutations were identified in 199 clonally related sweet orange accessions (n=19.085). Here these mutations are referred to as “clonal_ACCESSION_”. **(a)** The proportion of somatic mutations in genic regions relative to the genomic expectation, **(b)** dN/dS calculated as the ratio of nonsense to synonymous mutations relative to the genomic expectation, and **(c)** dN/dS calculated as the ratio of missense to synonymous mutations relative to the genomic expectation for the sets of leaf-specific and mosaic mutations detected within the historic tree and the set of clonal mutations detected in 199 clonal accessions. Points are colored by whether they are significantly different from the genomic expectation (red = enriched, blue = depleted) or not (black). The genomic expectation is calculated as the mean value for 1000 permutations of randomly introduced *in silica* mutations (gray points) of the same sample size and mutation spectrum as the focal mutation set. Mutations in subtelomeric regions were excluded. Raw ratios and p-values are detailed in Tables S5 and S7.

Reduction in genic mutations could result either from somatic purifying selection to remove deleterious mutations in genic sequences or from adaptive mutation bias to prevent the occurrence of genic mutations. There is evidence that adaptive mutation bias shapes mutation accumulation patterns in plants (40, 41), with epigenomic features such as reduced DNA methylation levels in protein-coding sequences associated with lower mutation rates (41). To distinguish between the effects of mutation bias and somatic selection, we focused on the accumulation of nonsynonymous somatic mutations (42). While mutation bias will reduce both nonsynonymous and synonymous mutations, somatic selection will result in differential accumulation of deleterious mutations (nonsynonymous) relative to silent (synonymous) mutations. We first focused on nonsense mutations because the introduction of a premature stop codon is most likely to have a deleterious effect. The ratio of nonsense to synonymous mutations was calculated for leaf-specific, mosaic, and clonal mutations relative to the genomic expectation based on permutations. No nonsense mutations were identified in the set of 218 clonal mutations (Table S7). A larger number of nonsynonymous mutations were identified across the set of clonally related accessions (nonsense = 99; missense = 1,537). Nonsense mutations were depleted in the sets of mosaic and clonal mutations relative to synonymous mutations, suggesting somatic purifying selection is shaping patterns of mutation accumulation in plant meristematic cells (mosaic: n = 2,725, 1000 permutations, p = 0.005; clonal: n = 19,085, 1000 permutations, p = 0.001) (Figure 4b, Table S7). Unlike the mosaic and clonal mutations, nonsense mutations were not depleted relative to synonymous mutations for leaf-specific mutations indicating somatic purifying selection is weaker or does not occur in differentiated cells (n = 1,906) (Figure 4b, Table S7).

Studies of somatic mutation in the human body have concluded that positive somatic selection drives patterns of somatic mutation accumulation and cellular evolution (20–22, 65). During tumorigenesis, driver mutations which increase cell fitness lead to clonal expansion of a cell lineage. In plants, however, the expansion of clonal cell lineages is not expected due to the rigid structure of the extracellular matrix (i.e. cell walls) (66). Surprisingly, we observe evidence of positive somatic selection in both leaf-specific and mosaic somatic mutations (leaf-specific: n = 1,906; mosaic: n = 2,725) (Figures 4c and S17, Table S7). Unlike nonsense mutations, the effects of missense mutations on protein function are difficult to predict. To further understand the mechanisms underlying the elevated proportion of missense mutations in leaf-specific and mosaic mutations, we tested for over-representation of Gene Ontology (GO) terms assigned to 441 genes with missense mutations from the total set of 12,025 mutations identified in the historic tree. Over-represented GO terms included those related to stem cell or meristem maintenance and DNA repair (Figure S18). This suggests that mutations in genes which are important for cell division and the maintenance of genome integrity are subjected to positive somatic selection, mirroring patterns of positive selection on driver mutations in cancer. There were also several enriched GO terms that were related to pathogen defense response (Figure S18). The hypothesis that somatic mutations may confer beneficial phenotypic mosaicism is central to the genetic mosaic theory of plant defense (67, 68). Intra-individual phenotypic variation in pest and herbicide resistance has been observed in *Populus*, *Eucalyptus*, and *Hydrilla* and is suspected to be caused by mosaic somatic mutations (69). Our results provide preliminary support for this phenomenon, but further investigation is needed to confirm the role of these mutations in the plant’s defense response.

## Discussion

The relevance of genetic mosaicism to human health has led to an abundance of research on understanding the processes which shape somatic mutation accumulation in humans. Despite the importance of genetic mosaicism to plants that reproduce asexually, including many crop species, the forces which shape the accumulation of somatic mutations in plants are comparatively unexplored. We have described the patterns and processes, including somatic selection, that shape somatic mutation in a single, long-lived navel orange tree and an associated clonal lineage derived from and maintained by asexual propagation.

A number of studies in long-lived or clonally propagated plant species have surveyed genome-wide patterns of somatic mutation accumulation (10–12, 24, 27, 29, 30, 50, 51, 70–74). Since a majority of somatic mutations are at low mutation fractions (Figure 2c-d) (11, 15, 30), somatic mutation detection in some of these studies has been limited by low depths of sequencing and methods for variant identification that do not consider low fraction mutations. Somatic mutations restricted to a specific meristematic stem cell layer will occur at low mutation fractions. Layer-specific somatic mutations have previously been identified in other plant species (51, 74) and we found good correspondence between the mutation fraction of somatic mutations detected in neighboring leaves (and branches), suggesting the fixation of layer-specific mutations is common (Figure S4). The fixation of mutations in specific layers, even if they are at low mutation fractions, is important to consider because these mutations can be maintained by asexual propagation (i.e. periclinal chimeras) and if they occur in a specific meristematic stem cell layer (L2) they can also be transmitted to sexual progeny (74). The transmission of low fraction mutations has been demonstrated for at least one tree species (30). Some citrus species are able to produce embryos from maternal tissue, also arising from the L2 meristematic layer, offering another path of transmission for somatic mutations (75).

The identification of genuine somatic mutations at low mutation fractions was essential for exploring the dynamics of mutations within a long-lived navel orange tree and across a related clonal lineage of sweet oranges. There was good evidence that somatic purifying selection shapes mutation accumulation, particularly for mosaic mutations within the historic tree and for clonal mutations that vary between clonally related sweet orange accessions. Evidence of somatic purifying selection has also been described previously, including in a species of tropical trees and a clonal sea grass (24, 30). However, others find that somatic mutations are selectively neutral (29). It is likely that the strength of somatic purifying selection can vary between organisms, but until practices for somatic mutation identification are unified it is difficult to compare across organisms and studies. A majority of new somatic mutations will be heterozygous (in the cell where they arise), so how does selection act on these new mutations? This is a major outstanding question. The effect of nonsynonymous mutations occurring in hemizygous regions will be instantly exposed to the effects of selection. Unfortunately, we did not identify a sufficient number of somatic mutations in hemizygous regions to evaluate whether the strength of somatic purifying selection was elevated in these regions. New nonsynonymous somatic mutations may also be exposed to selection if their effects are dominant or if the alternate allele has been previously inactivated. Further research is needed to disentangle these scenarios and to understand the basis of somatic purifying selection in individual trees and their clonal lineages.

There are overwhelming signatures of somatic positive selection in studies of tumorigenesis in humans (20–22). Unlike tumor cells, the rapid expansion of clonal cell lineages in plants is not expected due to the rigid structure of cell walls (66). We also find signatures of positive selection for somatic mutations occurring within the historic tree, but not between clonal accessions (Figure 4c). These mutations occur in genes that function in cell division and DNA repair, similar to driver mutations that rapidly expand in human tumor cell lineages (76). Interestingly, these driver-like mutations were identified for somatic mutations that occur in a single leaf, which arose after the most recent branching event, indicating that these mutations may expand quickly after they occur. Overall, the action of somatic selection in genetically mosaic individuals will affect the number and types of mutations transmitted to both asexual and sexual progeny. In both cases, somatic mutations, similar to mutations that arise in the gametes, will contribute to segregating genetic variation which is the substrate for evolution.

## Materials and methods

### Sample collection and sequencing for the historic “Parent Washington Navel Orange Tree”

Mature leaves were collected in January of 2022 from the Parent Washington Navel Orange Mother Tree (*Citrus sinensis* (L.) Osbeck) located at the corner of Magnolia Ave and Arlington Ave in Riverside, CA. The tree was 149 years old at the time of sampling. Two adjacent leaves were collected at the branch apex and branch base for each of seven main branches. DNA was isolated from leaf tissue collected from the leaf tip and avoided the midrib using the Qiagen DNeasy Plant kit. Libraries were prepared using the Illumina DNA Prep kit. Illumina sequencing was performed using the NovaSeq platform at the DNA Technologies and Expression Analysis Cores at the UC Davis Genome Center.

### Sample collection and sequencing for clonal sweet orange accessions

Leaf tissue was collected from 88 sweet orange trees located in the Givaudan Citrus Variety Collection at the University of California, Riverside (Tables S4 and S9). For Illumina sequencing, young leaves were collected from a single leaf or cluster of newly emerging leaves in the tree canopy. DNA was extracted from leaf tissue using a CTAB extraction protocol with the addition of a sorbitol wash step (77). Libraries were prepared using the Illumina DNA Prep kit. Illumina sequencing was performed using the NovaSeq platform at the DNA Technologies and Expression Analysis Cores at the UC Davis Genome Center. For PacBio sequencing, mature leaves were collected from multiple locations of the tree canopy and combined prior to DNA extraction. High molecular weight DNA was isolated using the protocol “Preparing Arabidopsis Genomic DNA for Size-Selected ∼20 kb SMRTbellTM Libraries” with the addition of a sorbitol wash step following nuclei isolation and followed by purification using AMPure PB beads. Library prep and sequencing was performed using the PacBio Sequel II platform at the DNA Technologies and Expression Analysis Cores at the UC Davis Genome Center.

### Illumina sequencing data processing and variant calling for the Parent Washington Navel Orange Tree samples

Raw reads in FASTQ format were converted to BAM format using GATK FastqToSam. Adapter sequences were marked using GATK MarkIlluminaAdapters. Reads were aligned to the ‘Washington navel’ reference genome (49) using bwa-mem2 (78). Duplicate reads were removed using Picard MarkDuplicates. Variants were called relative to the ‘Washington navel’ reference genome using GATK HaplotypeCaller with ploidy=4 for all samples, followed by joint calling with GATK GenotypeGVCFs with a minimum confidence threshold of 30. Variants were then filtered to remove sites with missing data in one or more samples, multi-allelic variants, and variants immediately adjacent to homopolymer runs of 3 bases or larger. Homopolymer runs in the reference genome were identified using the make_homopolymer function from the polymorphology2 package in R (79). Variants were annotated using SnpEff (80). The mutation fraction was calculated as the number of reads supporting the alternate allele over the total number of reads supporting the position (alternate + reference allele).

### Illumina sequencing data processing and variant calling for clonal accession samples

Raw reads in FASTQ format were converted to BAM format using GATK FastqToSam. Adapter sequences were marked using GATK MarkIlluminaAdapters. Reads were aligned to the ‘Washington navel’ reference genome (49) using bwa-mem2 (78), then sorted and indexed using samtools. Duplicate reads were removed using Picard MarkDuplicates. Variants were called relative to the ‘Washington navel’ reference genome using GATK HaplotypeCaller with ploidy=1 for all samples, followed by joint calling with GATK GenotypeGVCFs with a minimum confidence threshold of 30. Variants were filtered to remove sites with missing data in one or more samples, and multi-allelic variants. Variants were annotated using SnpEff (80).

### PacBio HiFi sequencing data processing and variant calling for clonal accessions samples

Reads were aligned to the ‘Washington navel’ reference genome (49) using pbmm2. SNVs were called using GATK HaplotypeCaller with a higher ploidy parameter (ploidy=10) to increase the sensitivity of mutation detection in these samples given the higher sequencing depth than the Illumina sequencing data. SNVs were filtered to remove sites with missing data in one or more samples and multi-allelic variants. SNVs were annotated using SnpEff (80). SVs were called using Sniffles2 both with and without the mosaic option (81).

### Calculating the mutation fraction

The mutation fraction (MF) was calculated as the number of reads supporting the alternate allele over the total number of reads supporting the variant position (alternate + reference allele). Because variants were called relative to a single haplotype of the diploid haplotype-resolved ‘Washington navel’ reference genome, the MF calculated here is used as a proxy for the cell fraction in which the mutation is present within the sampled tissue. A heterozygous mutation present which is fixed in all sampled cells will be detected at a MF of 1, while a heterozygous mutation present in only a portion of sampled cells will be detected at a MF less than 1.

### Simulating *in silico* somatic mutations

Illumina sequencing reads generated using a NovaSeq platform were simulated using InSilicoSeq v1.5.4 (82). Three sets of simulated sequencing reads were generated from the ‘Washington navel’ reference assembly:

1. Reads simulated at 40X per-haplotype sequencing depth across a single diploid chromosome (Scaffolds 1A and 1B).
2. Reads simulated at 40X per-haplotype sequencing depth across a single haploid chromosome (Scaffold 1A).
3. Reads simulated at 10X per-haplotype sequencing depth across all 18 chromosome-level scaffolds.

*In silico* mutations were introduced into the first two sets (at 40X depth) of simulated sequencing reads using Mutation-Simulator at MFs ranging from 0.1 to 1 (83). The third set of simulated sequencing reads (at 10X depth) was left unmutated to assess false positives. Simulated sequencing reads were aligned to the ‘Washington navel’ reference genome using bwa-mem2 (78). Variants were called using GATK HaplotypeCaller with a ploidy setting of 1-10 and filtered to retain only bi-allelic single nucleotide variants. The recall rate was calculated as the number of *in silico* mutations which were detected over the total number of *in silico* mutations introduced. The specificity rate was calculated as the number of *in silico* mutations which were detected over the total number of variants called.

### Evaluating mutation detection using a haplotype-resolved reference assembly

To assess the impact of using a haplotype-resolved diploid reference genome on somatic mutation detection, simulated sequencing reads from both haplotypes of one chromosome were aligned to a haploid reference genome representing only a single haplotype. The specificity rate was lower (2.0 - 2.2%) when mapping reads to a haploid reference genome, resulting in a larger number of false positives (Figure S3e). The recall rate was also reduced, though this difference was less extreme. Additionally, the called MF of successfully detected *in silico* somatic mutations did not accurately reflect the MF at which they were introduced (Figure S3b). *In silico* somatic mutations were often called at approximately half the introduced MF, presumably due to the presence of additional reads from the unrepresented haplotype being mapped to the region containing the mutation. To test whether these results were indeed due to the mis-mapping of reads derived from a haplotype unrepresented in the reference, we mapped only the subset of simulated sequencing reads which were generated from the represented haplotype to the haploid reference genome. This eliminated the large number of false positives and resulted in called MFs which matched well with the introduced MFs (Figure S3c,f). This indicates that the mis-mapping of reads originating from portions of a diploid genome which are unrepresented in the reference produces false positives and inaccurate MF calls, supporting the usefulness of a haplotype-resolved diploid reference genome for the accurate detection of somatic mutations (47, 48).

### Filtering to remove likely false positives

False positive mutations detected in simulated sequencing reads were examined to identify the distinguishing characteristics of false positives compared to genuine somatic mutations. A number of false positives occur due to the mapping of reads originating from genomic regions not represented in the reference assembly. These false positives were most often identified in all simulated samples and were called at MFs of approximately 0.5 (Figure S3e). Additionally, a smaller set of false positives were identified in the set of called variants when mapping reads to the full diploid reference genome. These false positives were always identified in only one of four simulated samples, and were called at low MFs (Figure S3d). Because the set of these false positives was small (n=185), we further generated three sets of 10X (per-haplotype) simulated sequencing reads from the entire diploid reference genome. These reads were left unmutated and mapped to the diploid reference. Variants were called using GATK HaplotypeCaller with a ploidy parameter setting of 4. There were an average of 1,363 false positives called in each simulated sample. The vast majority of these false positives were specific to only one simulated sample and the alternate alleles of the false positive variants were supported by only 1-2 reads. In conclusion, there appears to be two categories of false positive variants when attempting to discover genuine somatic mutations. The first category originates from reads which derive from genomic regions which are unrepresented in the reference assembly, and is characterized by variants called in all samples at MFs near 0.5. The second category of false positives originates from sample-specific sources of error and is characterized by variants called in a single sample and supported by few reads. Variants detected in all samples were removed as likely false positives if the called MF was between 0.4 and 0.6 (n = 144). Variants were additionally removed if the site was supported, on average across, by fewer than 10 or more than 100 reads per sample (n = 136,560).

### Phylogenetic trees

Distance matrices were constructed from somatic mutation MF matrices using the dist function (Euclidean method) in R (84), which were then used to construct phylogenetic trees using the nj function from the ape package in R (85). Trees were visualized using the ggtree package in R (86).

### Mutation spectra

The mutation spectra was calculated as the proportion of mutations of each base change in a trinucleotide context relative to the frequency of each respective trimer in the reference genome. Trimer frequencies were determined using the genome_Nmer_frequencies function from the polymorphology2 package in R (79). Mutation spectra were plotted using a modified version of the plot_tricontexts function from the polymorphology2 package in R (79).

### Subtelomeres

Tandem repeats arrays were identified in the ‘Washington navel’ reference genome using Tandem Repeats Finder (Version 4.09, 2020) (87). The subtelomeres were defined as tandem repeat arrays containing the 181bp subtelomeric repeat sequence.

### Calculating the proportion of genic and nonsynonymous mutations relative to the genomic expectation

The proportion of genic and nonsynonymous:synonymous mutations was calculated and compared to the genomic expectation. For each data set the genomic expectation was calculated as the mean proportion of nonsynonymous:synonymous mutations among 1000 sets of randomly introduced *in silico* mutations with the same sample size and mutation spectrum as the observed mutation set.

### GO term enrichment

GO terms were assigned to genes using OMA (88). Over-represented GO terms were selected as those which were assigned to a significantly higher proportion of genes in the set of genes containing missense mutations relative to all genes (Fisher’s exact p<0.01).

## Acknowledgements and funding sources

We thank the Vidalakis group and Sohrab Bodaghi for providing access to the historic “Parent Washington Navel Orange Tree”. This research was supported by UCR initial complement funds and funding from the Citrus Research Board to D.K.S (Project 5200-201E); T.R.B is a fellow in the Plants-3D NSF National Research Traineeship Program (DBI-1922642). E.A.D was supported by a UC Mexus Postdoctoral Fellowship. Computations were performed using the computer clusters and data storage resources of the UC Riverside HPCC, which were funded by grants from NSF (MRI-2215705, MRI-1429826) and NIH (1S10OD016290-01A1). Sequencing of Illumina and PacBio libraries was performed by the DNA Technologies and Expression Analysis Cores at the UC Davis Genome Center, supported by NIH Shared Instrumentation Grant 1S10OD010786-01.

## Author contributions

T.R.B and D.K.S designed the study. T.R.B performed the analysis with support from E.A.D. E.A.D produced and annotated the ‘Washington navel’ genome. T.R.B and D.K.S wrote the initial manuscript and all authors commented and contributed to the final manuscript.

## Data Availability

Raw sequencing reads from the PWN tree canopy are available under NCBI BioProject PRJNA1137877. Raw Illumina sequencing reads for the GCVC sweet orange accessions are available under NCBI BioProject PRJNA1272085. Raw PacBio sequencing reads for the GCVC sweet orange accessions will also be made available at NCBI (BioProject pending).

