## Supplemental Figures and Tables for "Somatic selection shapes mutation accumulation in *Citrus sinensis*"

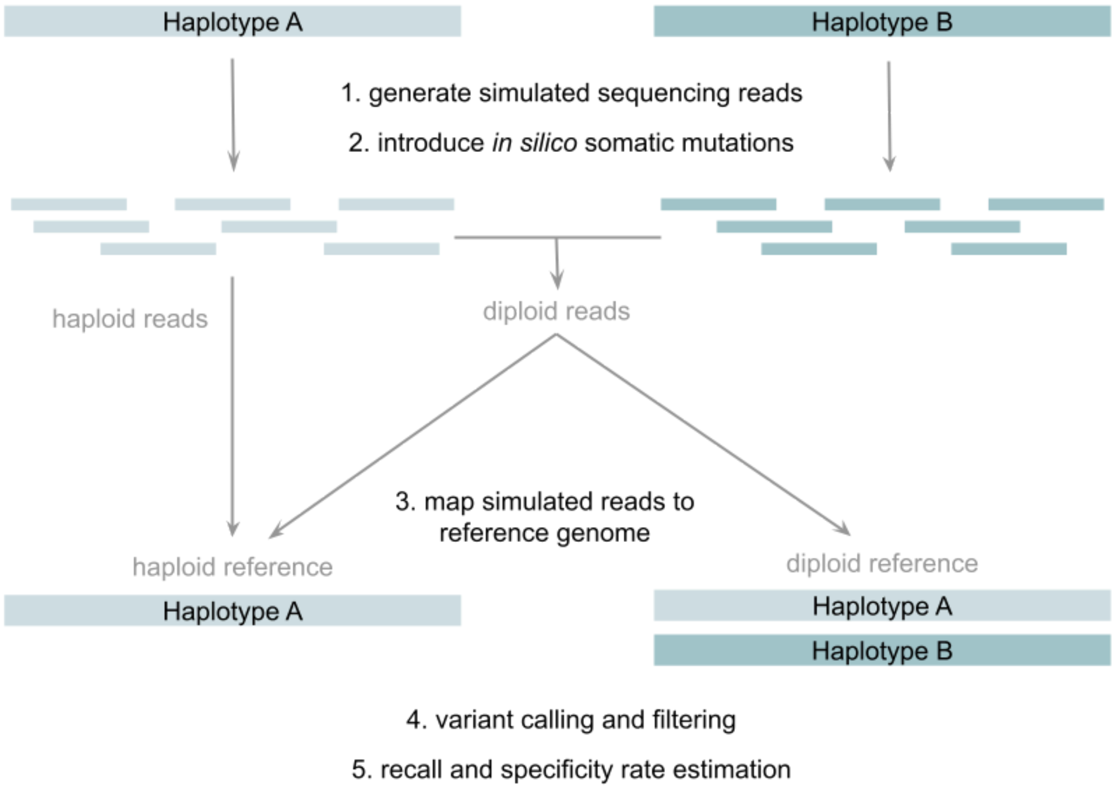


**Figure S1.** *In silico* somatic mutation analysis workflow.

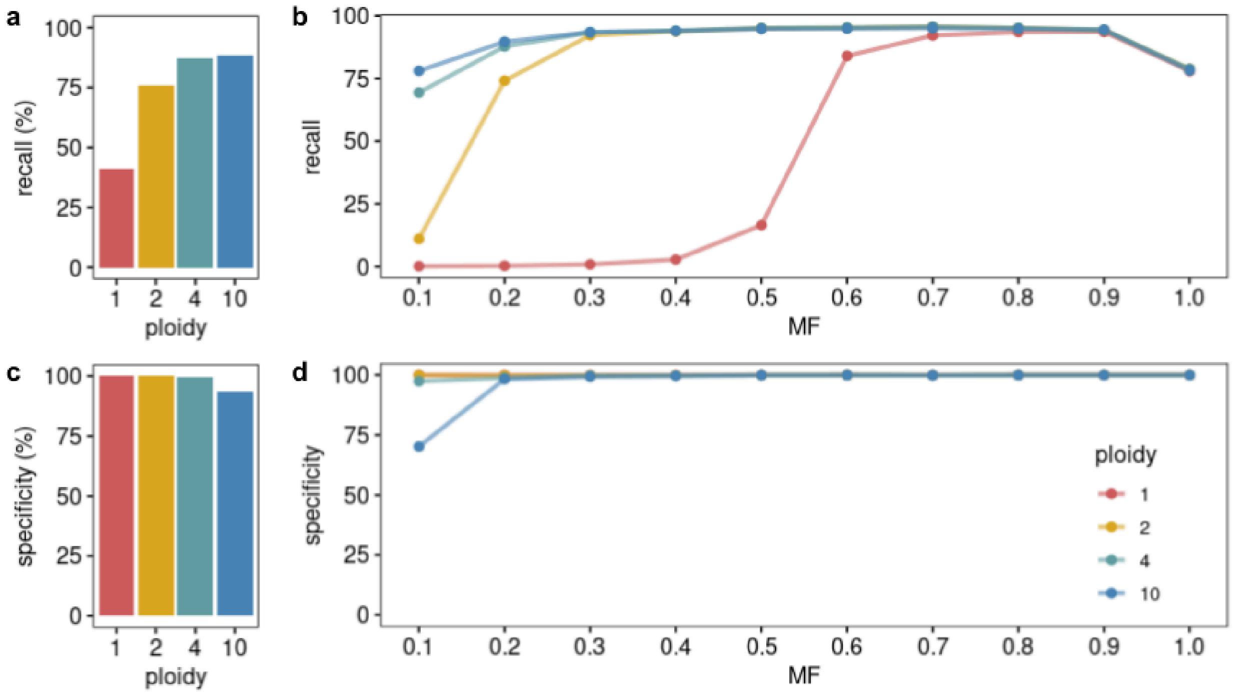


**Figure S2. *In silico* somatic mutation detection rates. (a-b)** The effect of increasing the GATK HaplotypeCaller ploidy parameter on the rate of recall for **(a)** overall *in silico* mutation detection and for **(b)** *in silico* mutation detection for MFs ranging from 0.1 to 1. The recall rate is the percentage of introduced *in silico* mutations which were called as variants. **(c-d)** The effect of increasing the GATK HaplotypeCaller ploidy parameter on the rate of specificity for **(c)** overall *in silico* mutation detection for overall and for **(d)** *in silico* mutation detection for MFs ranging from 0.1 to 1. The specificity rate is the percentage of variants called which were introduced *in silico* mutations.

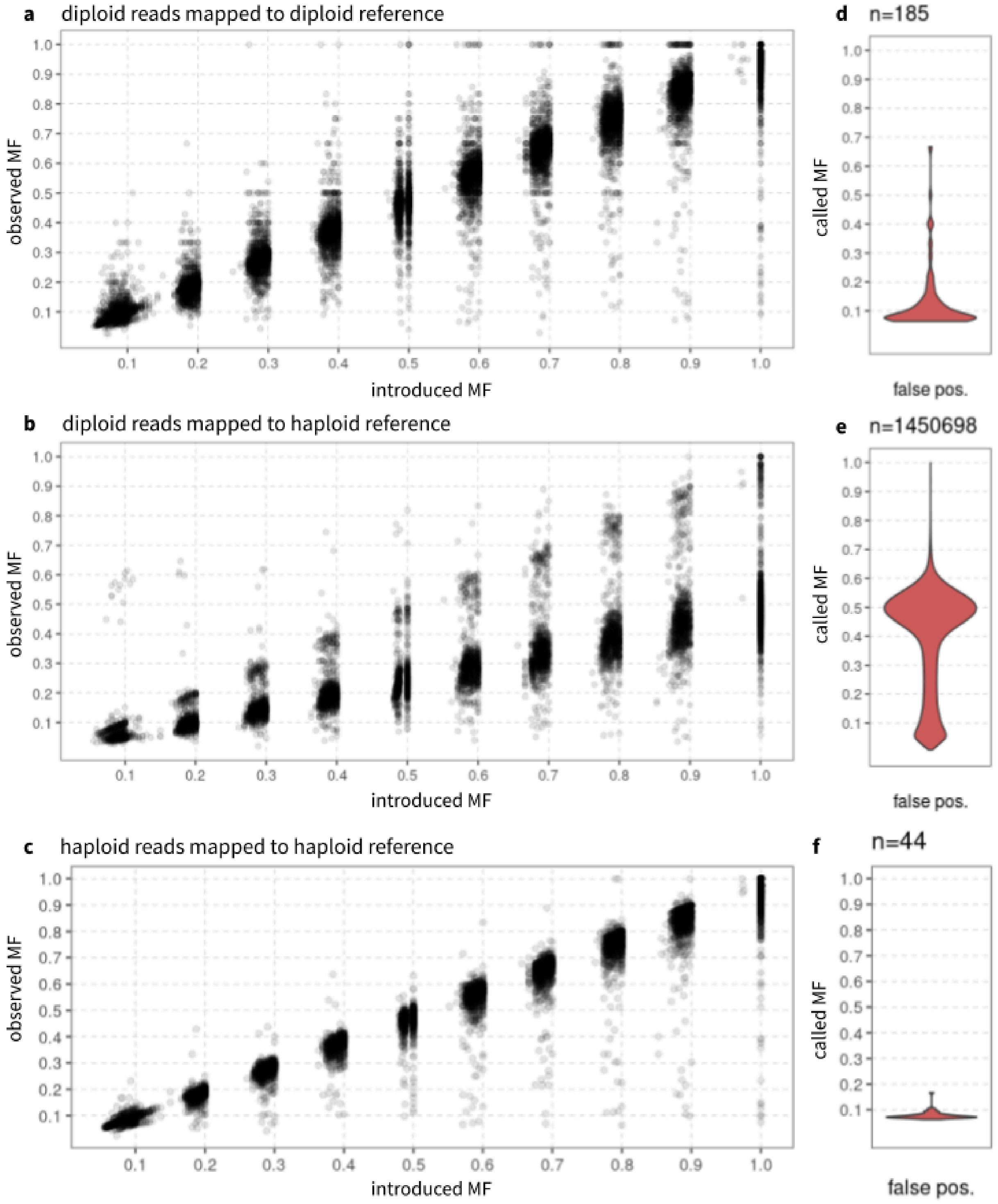

a diploid reads mapped to diploid reference
observed MF
introduced MF
d n=185
called MF
false pos.
b diploid reads mapped to haploid reference
observed MF
introduced MF
e n=1450698
called MF
false pos.
c haploid reads mapped to haploid reference
observed MF
introduced MF
f n=44
called MF
false pos.

**Figure S3. The MFs of *in silico* somatic mutations. (a-c)** The relationship between the introduced MF and the observed MF of *in silico* somatic mutations. Variants were identified using GATK HaplotypeCaller (with a ploidy parameter of 4) from **(a)** simulated reads from a diploid genome which were mapped to a diploid reference genome, **(b)** simulated reads from a diploid genome which were mapped to a haploid reference genome, and **(c)** simulated reads from a haploid genome which were mapped to a haploid reference genome. **(d-f)** The distribution of MFs of false positives identified using GATK HaplotypeCaller with a ploidy parameter of 4 from **(d)** simulated reads from a diploid genome which were mapped to a diploid reference genome, **(e)** simulated reads from a diploid genome which were mapped to a haploid reference genome, and **(f)** simulated reads from a haploid genome which were mapped to a haploid reference genome. The diploid reference genome refers to the ‘Washington navel’ genome assembly and the haploid reference genome refers to haplotype 1 of the same assembly.

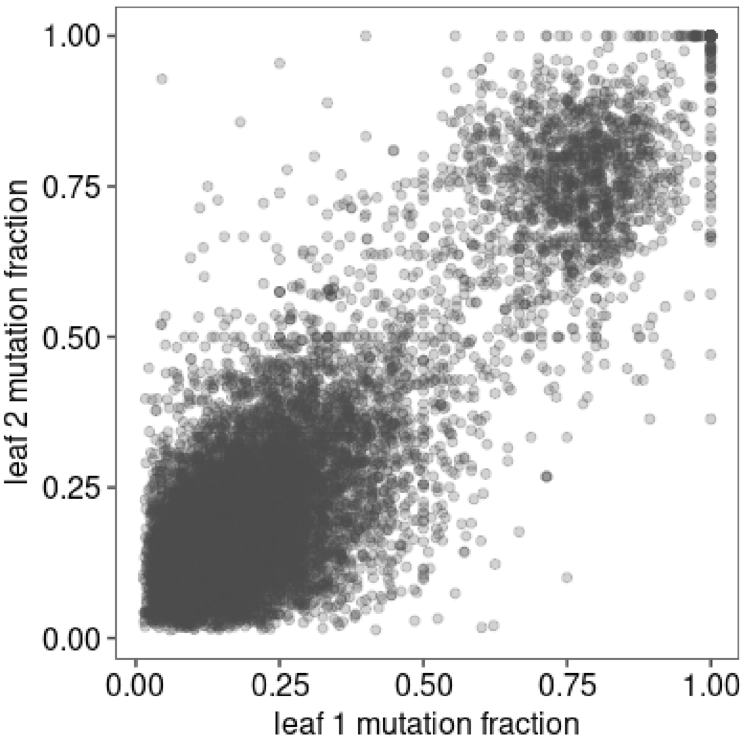


**Figure S4. Concordance between the observed mutation fraction at adjacent leaves.** Each point represents a mutation identified in adjacent leaves of a leaf-pair collected at one sampling location.

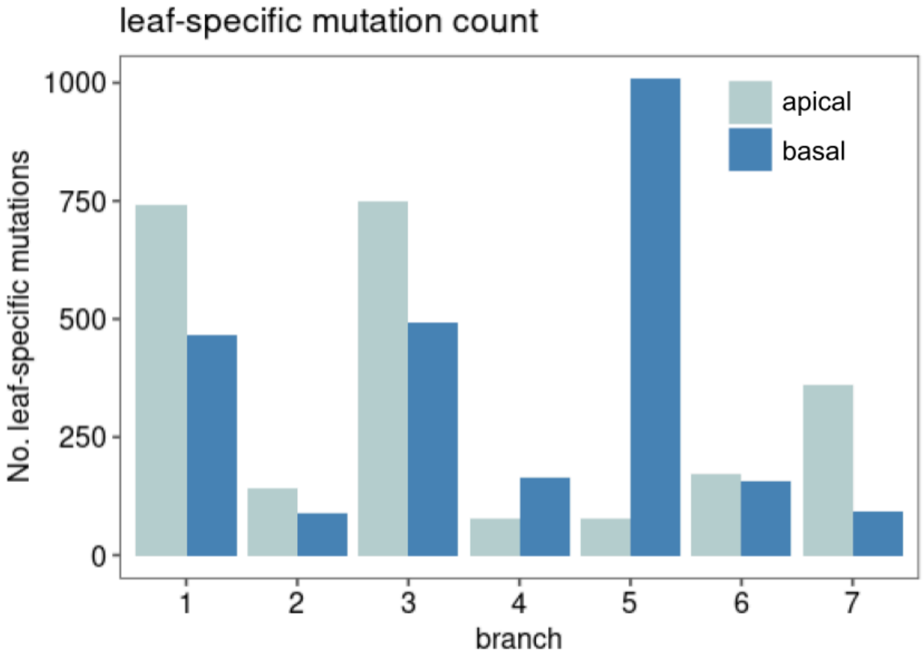


**Figure S5.** Number of leaf-specific somatic mutations detected at each sampling location.

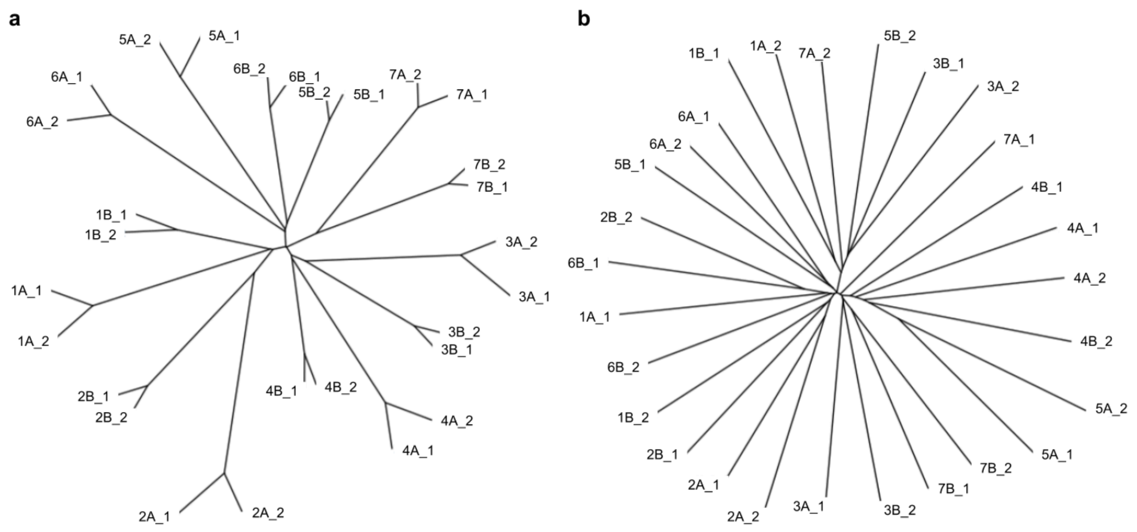


**Figure S6.** The phylogenetic trees constructed using the mutation fraction (MF) matrix of **(a)** completely-detected or **(b)** incompletely-detected mosaic somatic mutations.

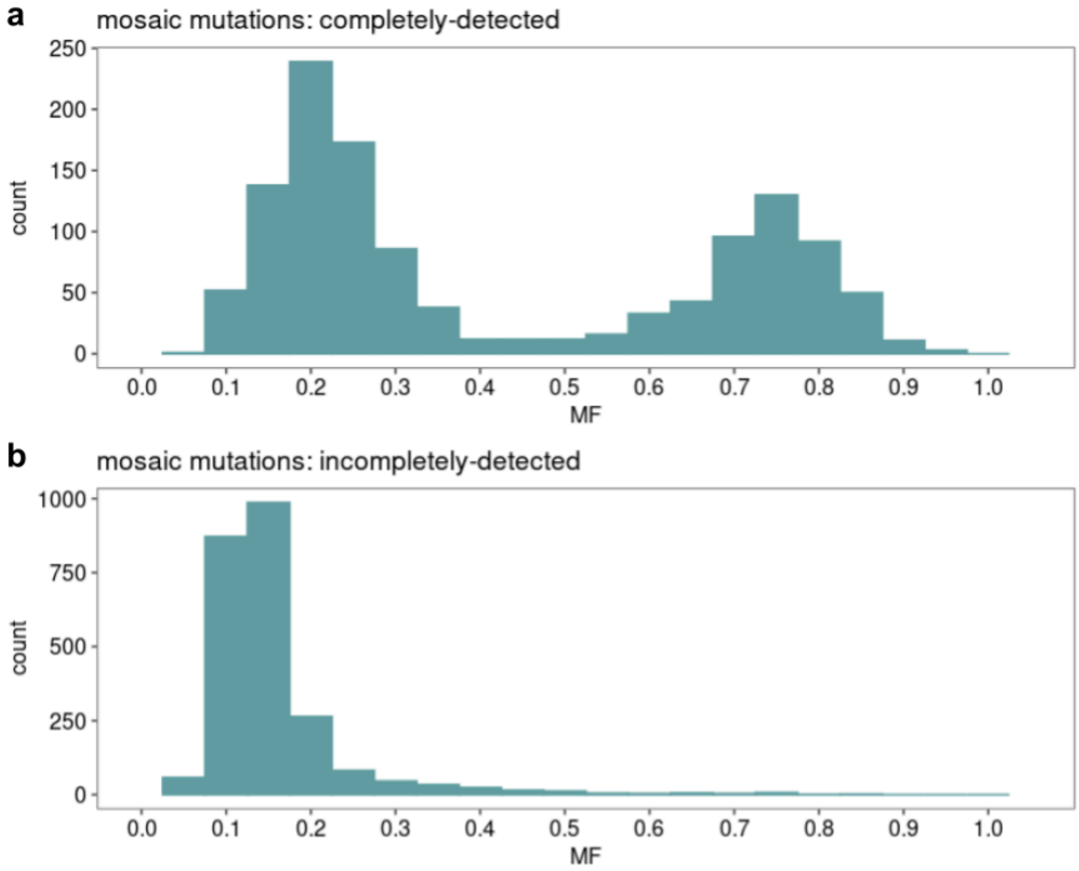


**Figure S7.** Mutation fraction (MF) distribution of **(a)** completely-detected and **(b)** incompletely-detected mosaic mutations.

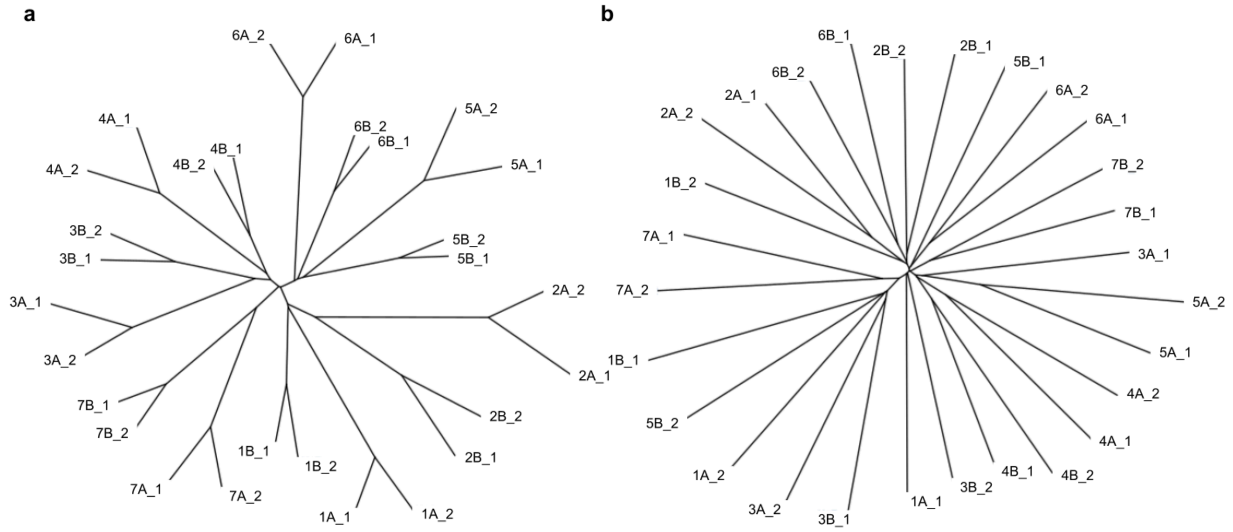


**Figure S8.** The phylogenetic trees constructed using the MF matrix of **(a)** high-fraction (MF > 0.5) or **(b)** low-fraction (MF < 0.5) mosaic somatic mutations.

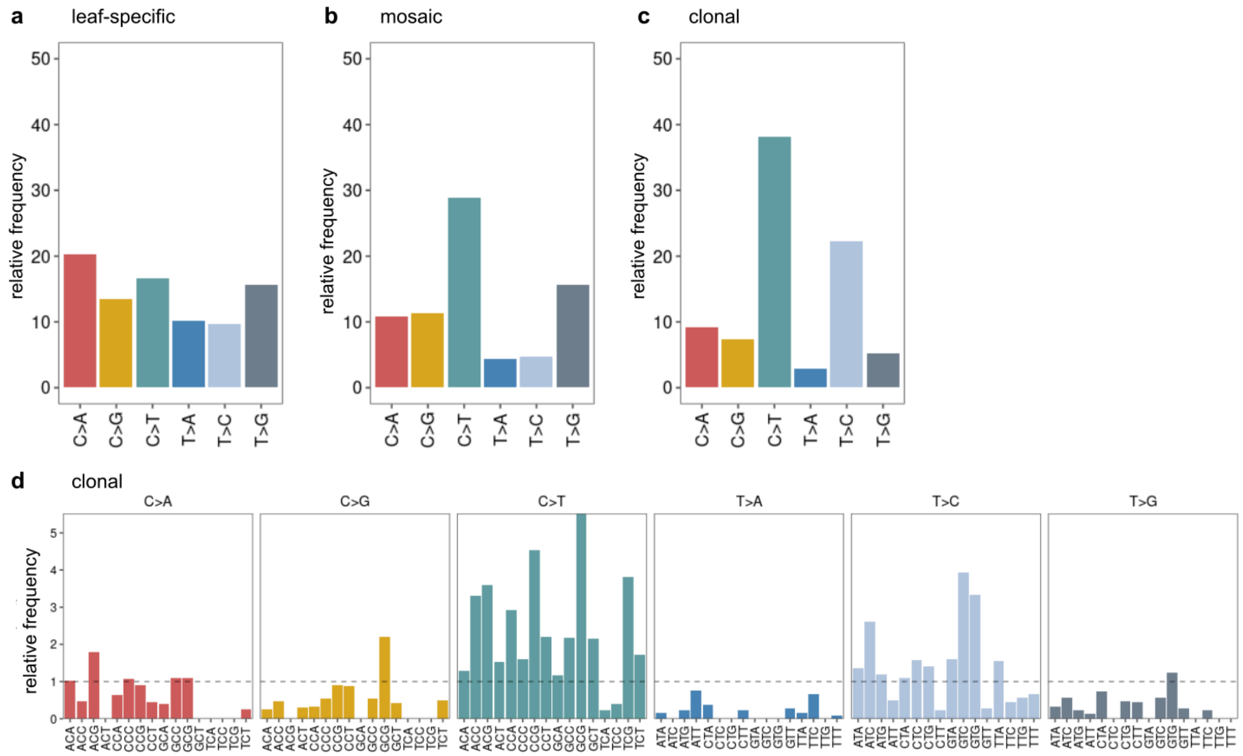


**Figure S9. Mutation spectra of leaf-specific, mosaic, and clonal somatic mutations. (a-c)** The sum of the proportion of each mutation class relative to the genomic trimer frequency for **(a)** leaf-specific, **(b)** mosaic, and **(c)** clonal somatic mutations in the historic tree. **(d)** The proportion of each mutation class relative to the genomic trimer frequency for the set of clonal somatic mutations for the set of clonal mutations identified in the historic tree (n=218).

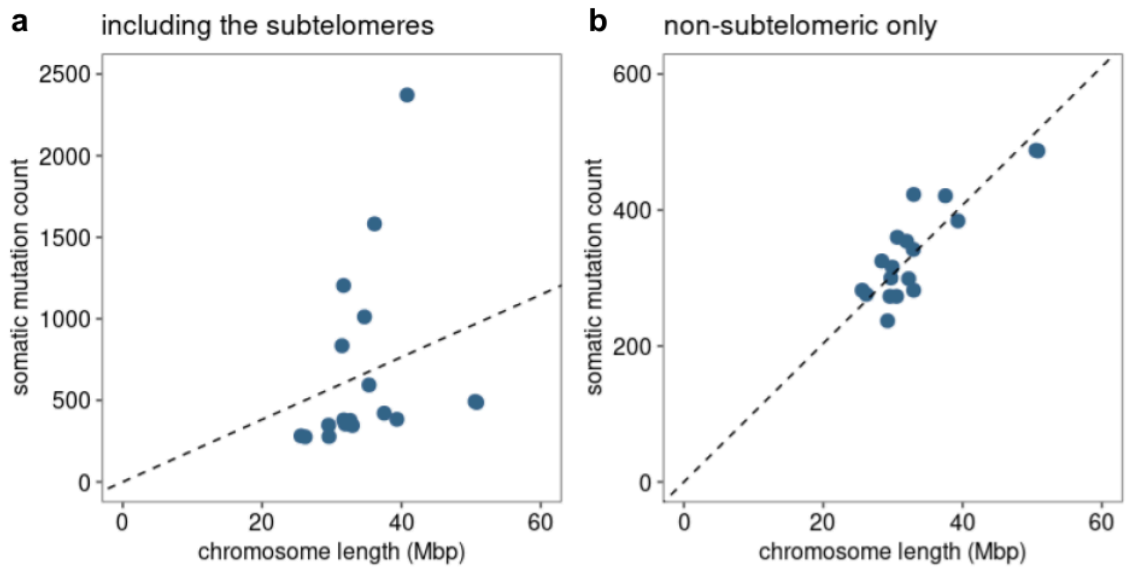


**Figure S10. The relationship between somatic mutation count and chromosome length. (a)** The number of somatic mutations detected per chromosome is not proportional to chromosome length. **(b)** The number of somatic mutations detected per chromosome is proportional to chromosome length when masking the subtelomeres.

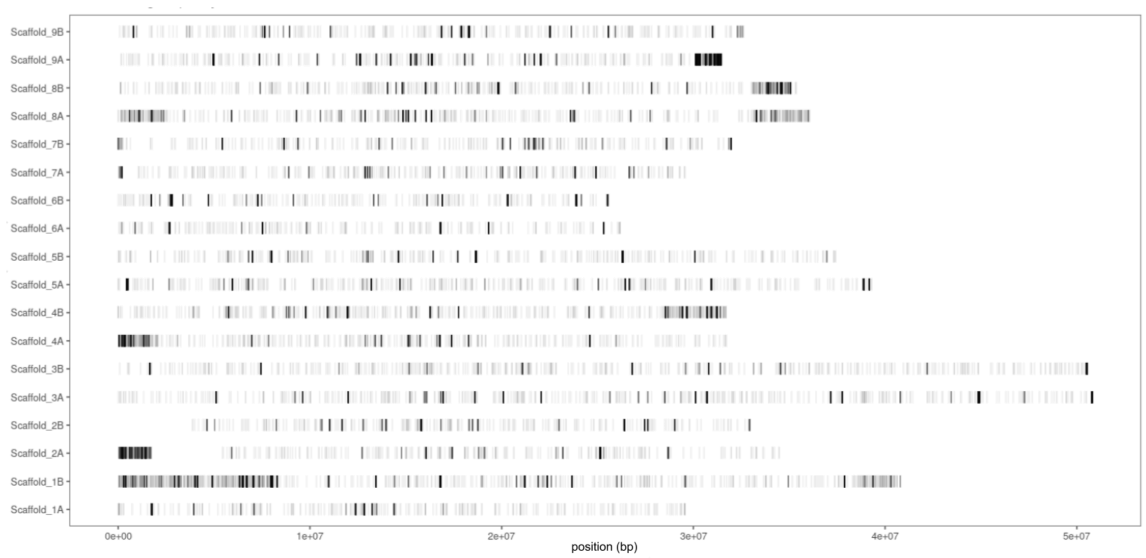


**Figure S11.** Chromosomal distribution of SNVs detected in 6 sweet orange individuals using PacBio Hifi sequencing data.

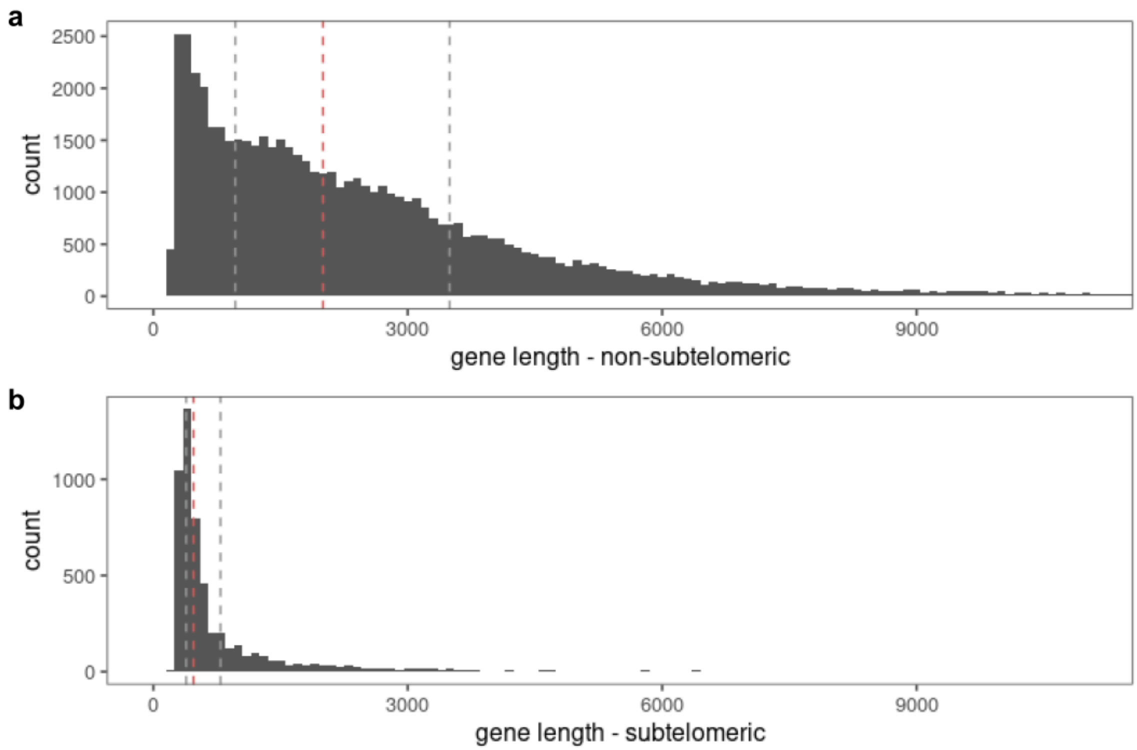


**Figure S12. (a-b)** Distribution of annotated gene length **(a)** in non-subtelomeric and **(b)** in subtelomeric regions. Red dashed lines indicate the mean gene length. Gray dashed lines indicate the first and third quartile

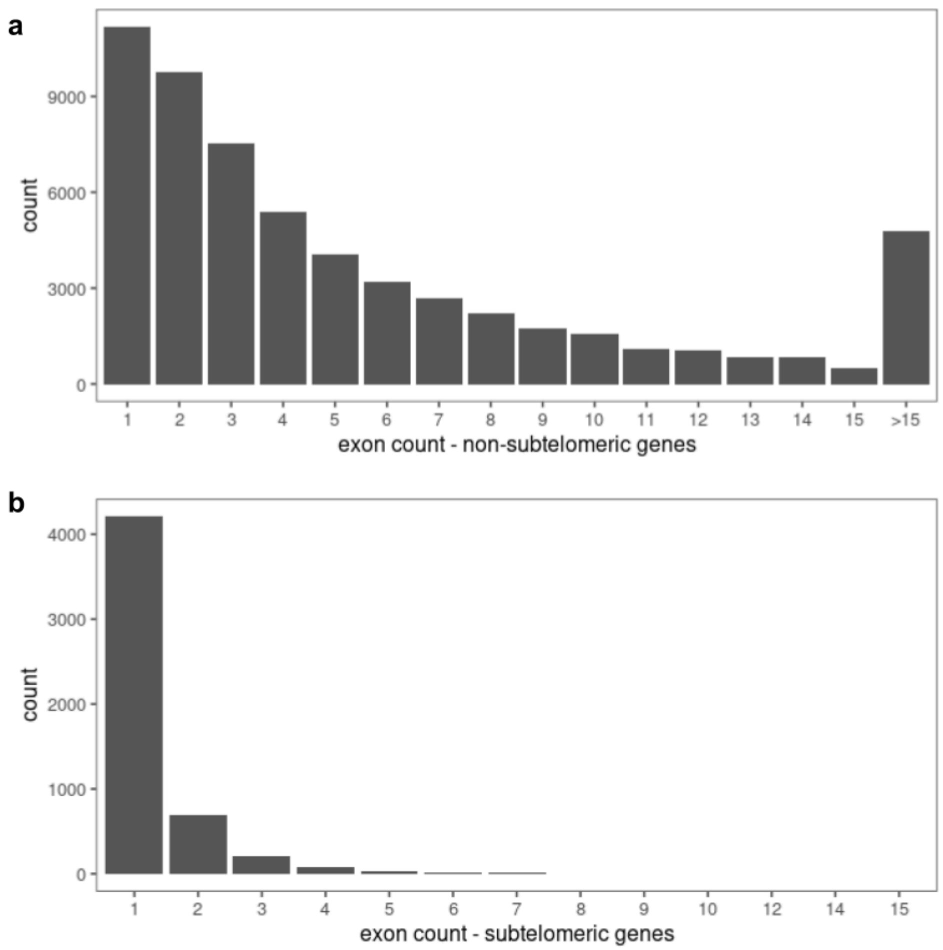


**Figure S13.** Number of exons in genes annotated **(a)** in the non-subtelomeric and **(b)** subtelomeric regions.

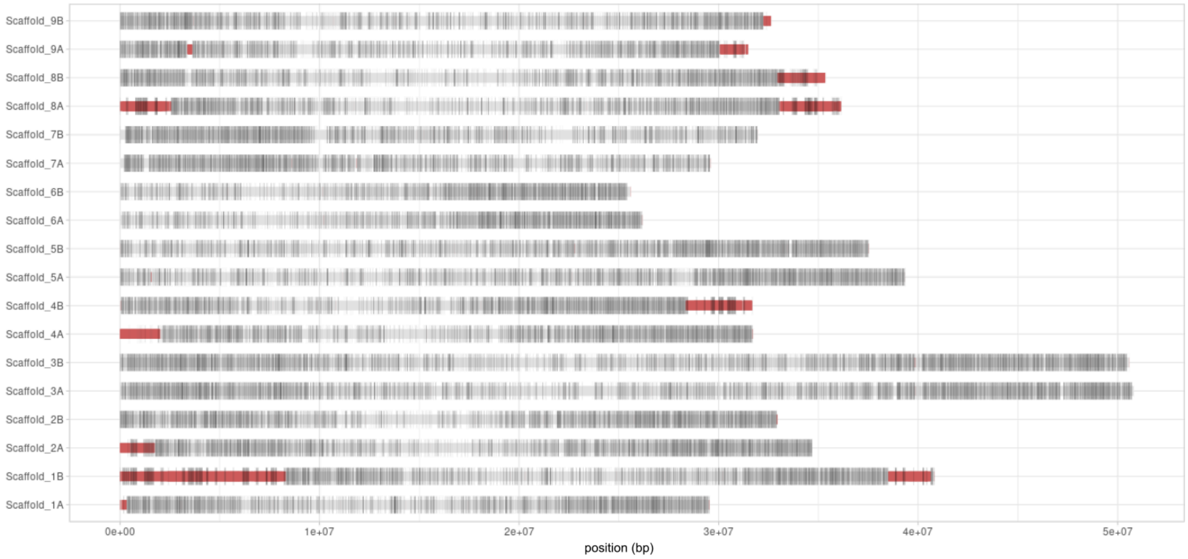


**Figure S14.** Genomic distribution of annotated protein-coding genes in the 'Washington navel' reference assembly. Subtelomeric regions are shown in red.

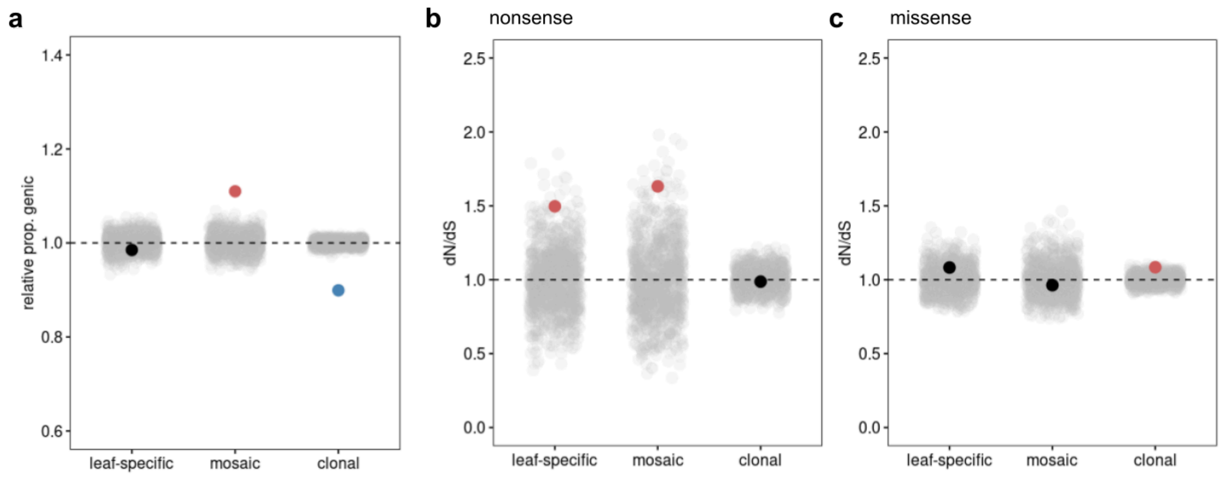


**Figure S15. Proportion of genic mutations and dN/dS in the subtelomeres**. Only mutations residing in subtelomeric regions were considered ($n_{leaf\text{-}specific}$ = 2888, $n_{mosaic}$ = 935, $n_{clonal}$ = 50). **(a)** The proportion of somatic mutations in genic regions relative to the genomic expectation based on permutation. **(b)** dN/dS calculated as the ratio of nonsense to synonymous mutations relative to the genomic expectation. **(c)** dN/dS calculated as the ratio of missense to synonymous mutations relative to the genomic expectation. Points are colored by whether they are significantly different from the genomic expectation (red = enriched, blue = depleted) or not (black). The genomic expectation is calculated as the mean value from 1000 permutations of randomly introduced *in silico* mutations (gray points) of the same sample size and mutation spectrum as the mutation set.

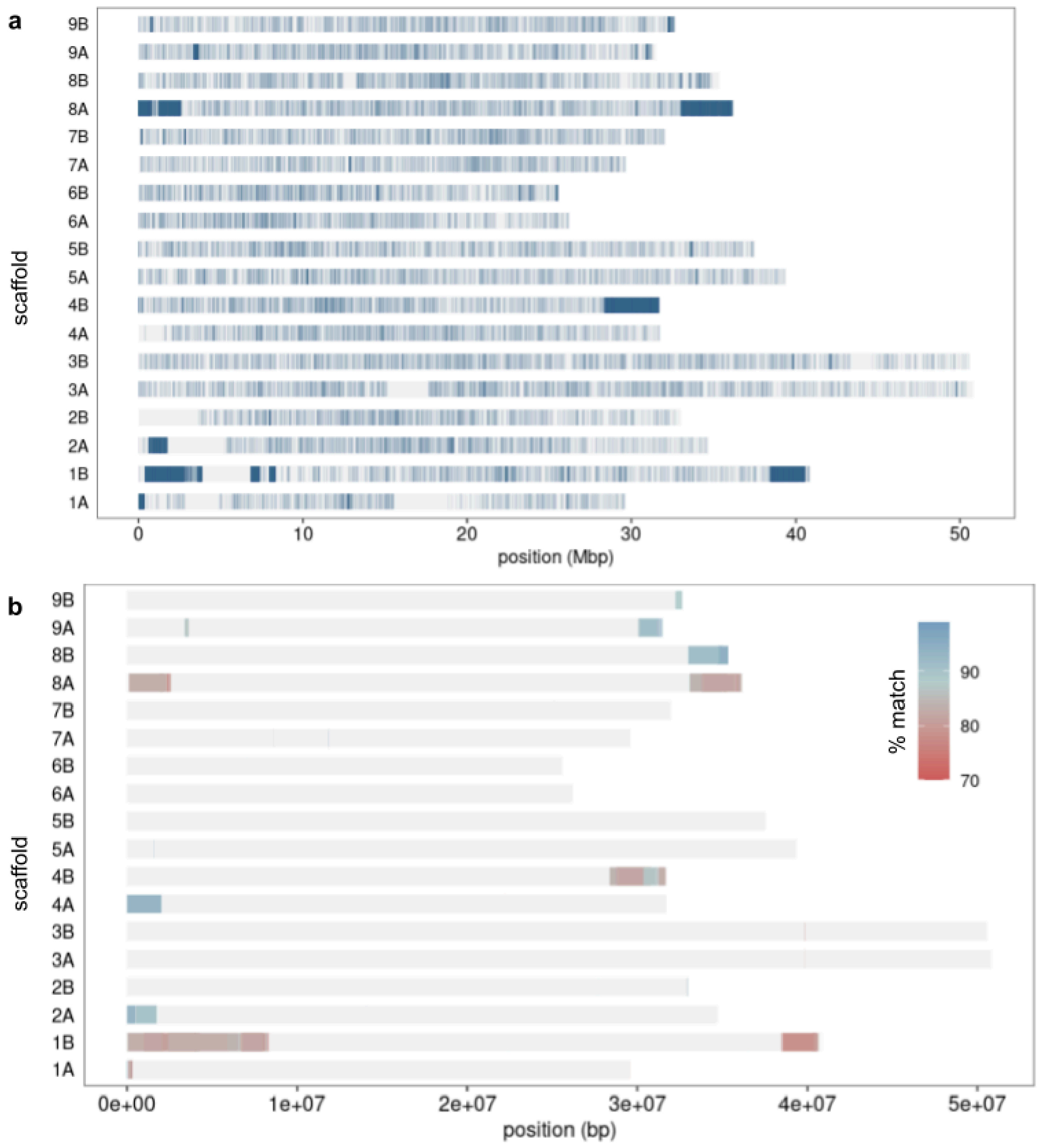


**Figure S16. Clonal mutation accumulation in the subtelomeres. (a)** The chromosomal distribution of SNVs detected in 199 clonally related accessions of sweet orange sequenced using Illumina short reads. **(b)** Subtelomeric regions colored by percent matches (percent of matches between adjacent copies overall).

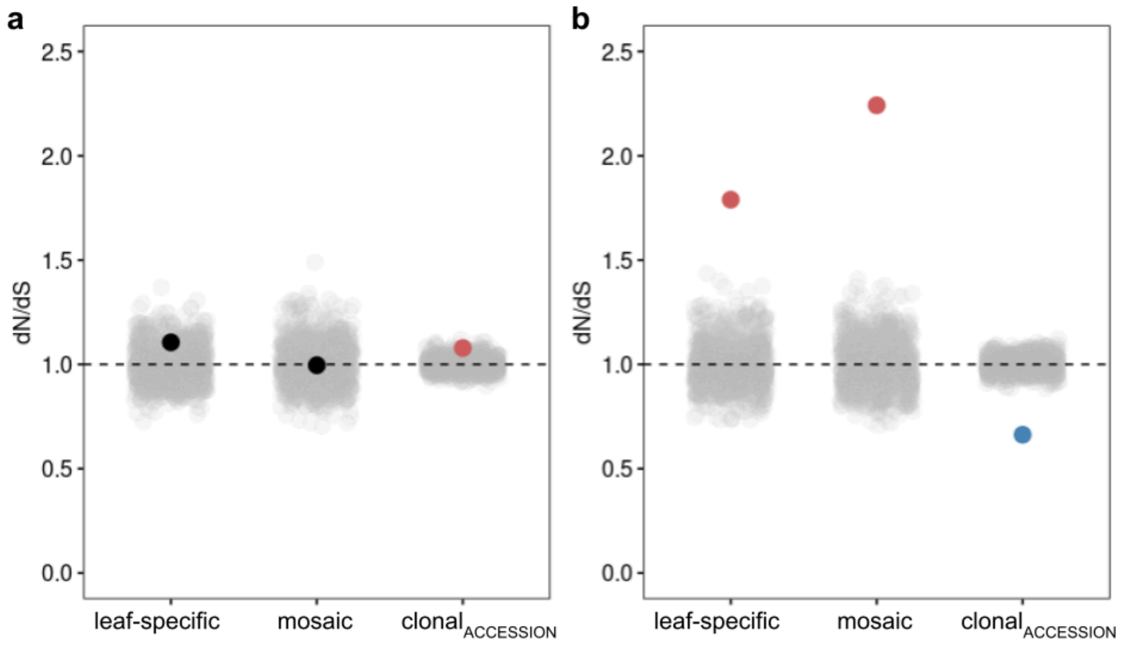


**Figure S17.** Combined dN/dS calculated as the ratio of nonsynonymous (nonsense + missense) mutations to synonymous mutations for the set of leaf-specific, mosaic, and clonal$_{ACCESSION}$ mutations **(a)** within the subtelomeres, and **(b)** outside of the subtelomeres. Points are colored if they are significantly different from the genomic expectation (red = significantly greater, blue = significantly lower) or black if they are not significantly different. The genomic expectation is calculated as the mean value from 1000 sets of randomly introduced *in silico* mutations with the same sample size and mutation spectrum as the mutation set (shown in gray).

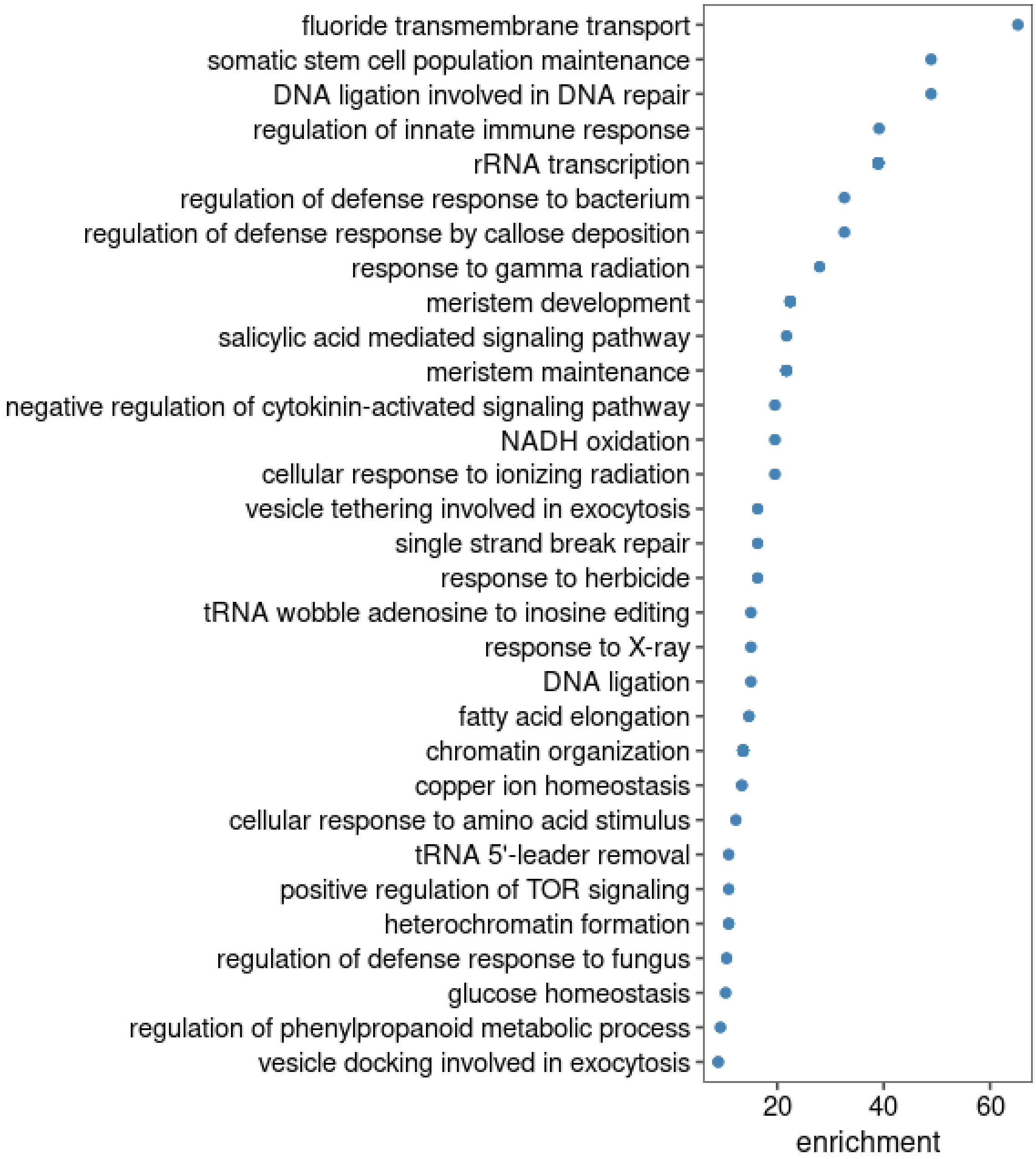


**Figure S18.** Biological process terms in the top 20% of over-represented GO terms among genes containing at least two missense mutations.

### Tables

**Table S1.** Sequencing & read mapping metrics. * Depth = per-haplotype average sequencing depth.

| Sample ID | Branch | Location | Leaf | Depth* | NCBI accession |
|---|---|---|---|---|---|
| 1A_1 | 1 | apical | 1 | 42.66 | SAMN42644081 |
| 1A_2 | 1 | apical | 2 | 42.21 | SAMN42644082 |
| 1B_1 | 1 | basal | 1 | 33.89 | SAMN42644083 |
| 1B_2 | 1 | basal | 2 | 48.86 | SAMN42644084 |
| 2A_1 | 2 | apical | 1 | 39.22 | SAMN42644085 |
| 2A_2 | 2 | apical | 2 | 43.91 | SAMN42644086 |
| 2B_1 | 2 | basal | 1 | 39.03 | SAMN42644087 |
| 2B_2 | 2 | basal | 2 | 32.26 | SAMN42644088 |
| 3A_1 | 3 | apical | 1 | 41.97 | SAMN42644089 |
| 3A_2 | 3 | apical | 2 | 46.31 | SAMN42644090 |
| 3B_1 | 3 | basal | 1 | 46.71 | SAMN42644091 |
| 3B_2 | 3 | basal | 2 | 45.15 | SAMN42644092 |
| 4A_1 | 4 | apical | 1 | 37.38 | SAMN42644093 |
| 4A_2 | 4 | apical | 2 | 37.35 | SAMN42644094 |
| 4B_1 | 4 | basal | 1 | 60.83 | SAMN42644095 |
| 4B_2 | 4 | basal | 2 | 35.05 | SAMN42644096 |
| 5A_1 | 5 | apical | 1 | 32.89 | SAMN42644097 |
| 5A_2 | 5 | apical | 2 | 34.83 | SAMN42644098 |
| 5B_1 | 5 | basal | 1 | 35.97 | SAMN42644099 |
| 5B_2 | 5 | basal | 2 | 46.13 | SAMN42644100 |
| 6A_1 | 6 | apical | 1 | 43.01 | SAMN42644101 |
| 6A_2 | 6 | apical | 2 | 42.05 | SAMN42644102 |
| 6B_1 | 6 | basal | 1 | 16.43 | SAMN42644103 |
| 6B_2 | 6 | basal | 2 | 41.59 | SAMN42644104 |
| 7A_1 | 7 | apical | 1 | 43.17 | SAMN42644105 |
| 7A_2 | 7 | apical | 2 | 36.79 | SAMN42644106 |
| 7B_1 | 7 | basal | 1 | 46.08 | SAMN42644107 |
| 7B_2 | 7 | basal | 2 | 43.84 | SAMN42644108 |

**Table S2.** Detection rates of *in silico* mutations introduced into 40X simulated sequencing data. Simulated sequencing reads were generated from a single chromosome pair of the diploid reference genome. * ploidy = GATK HaplotypeCaller ploidy parameter

| ploidy* | | introduced mutation fraction | | | | | | | | | |
|---|---|---|---|---|---|---|---|---|---|---|---|
| | | 0.1 | 0.2 | 0.3 | 0.4 | 0.5 | 0.6 | 0.7 | 0.8 | 0.9 | 1 |
| recall (%) | 1 | 0.1 | 0.3 | 0.8 | 2.8 | 16.5 | 84.0 | 92.2 | 93.5 | 93.6 | 77.9 |
| | 2 | 11.0 | 74.1 | 92.3 | 93.8 | 95.0 | 95.2 | 95.6 | 95.1 | 94.4 | 78.8 |
| | 4 | 69.4 | 87.8 | 93.3 | 94.1 | 95.2 | 95.4 | 95.7 | 95.2 | 94.5 | 78.8 |
| | 10 | 78.0 | 89.6 | 93.5 | 93.9 | 94.7 | 94.8 | 95.0 | 94.8 | 94.3 | 78.2 |
| specificity (%) | 1 | 100.0 | 100.0 | 100.0 | 100.0 | 100.0 | 100.0 | 100.0 | 100.0 | 100.0 | 100.0 |
| | 2 | 100.0 | 99.9 | 99.9 | 99.9 | 100.0 | 100.0 | 99.9 | 100.0 | 100.0 | 100.0 |
| | 4 | 97.5 | 98.8 | 99.7 | 99.8 | 100.0 | 100.0 | 99.9 | 100.0 | 100.0 | 100.0 |
| | 10 | 70.2 | 98.3 | 99.3 | 99.7 | 100.0 | 100.0 | 99.9 | 100.0 | 100.0 | 100.0 |

**Table S3.** Mutation density (number of mutations detected per Mbp) in subtelomeric and non-subtelomeric regions.

| | mutation density (per Mbp) | |
|---|---|---|
| | subtelomeric | non-subtelomeric |
| leaf-specific | 104.8 | 3.2 |
| mosaic | 33.9 | 4.5 |
| Illumina $clonal_{HISTORIC}$ | 1.8 | 0.3 |
| Illumina $clonal_{ACCESSION}$ | 323.8 | 31.8 |
| PacBio HiFi $clonal_{ACCESSION}$ | 176.5 | 34.8 |

**Table S4.** PacBio HiFi sequencing & read mapping metrics for samples collected from six sweet orange varieties. CRC number used to identify each accession in the Givaudan Citrus Variety Collection at the University of California, Riverside. Depth is calculated as per-haplotype average sequencing depth from reads mapped to the 18 primary scaffolds.

| Sample ID | CRC | Variety Name | Depth |
|---|---|---|---|
| PWN | 1241 | Parent Washington Navel | 40.4 |
| OLI | 2750 | Olinda Valencia | 35.5 |
| MOR | 3830 | Moro Blood orange | 41.9 |
| POW | 4040 | Powell Navel orange | 41.9 |
| CAR | 3994 | Cara Cara Navel orange | 42.3 |
| SHA | 4258 | Shahani Red Navel | 37.7 |

**Table S5.** Proportion of mutations in annotated gene regions ($prop_{mut}$) excluding the subtelomeres. The genomic expectation ($prop_{exp}$) is calculated as the mean of 1000 permutations. For $prop_{mut}$ values outside of a 95% CI, the p-value is the likelihood of a permutation value at least as extreme as the mutation value.

| | n | $prop_{mut}$ | $prop_{exp}$ | p-value |
|---|---|---|---|---|
| leaf-specific | 1906 | 0.661 | 0.682 | 0.001 |
| mosaic | 2725 | 0.641 | 0.682 | 0.001 |
| $clonal_{HISTORIC}$ | 168 | 0.554 | 0.680 | 0.001 |
| $clonal_{ACCESSION}$ | 19085 | 0.539 | 0.680 | 0.001 |

**Table S6.** Proportion of mutations in annotated gene regions ($prop_{mut}$) in the subtelomeres. The genomic expectation ($prop_{exp}$) is calculated as the mean of 1000 permutations. For $prop_{mut}$ values outside of a 95% CI of $prop_{exp}$, the p-value is the likelihood of a permutation value at least as extreme as the mutation value.

| | n | $prop_{mut}$ | $prop_{exp}$ | p-value |
|---|---|---|---|---|
| leaf-specific | 2888 | 0.377 | 0.383 | -- |
| mosaic | 935 | 0.426 | 0.383 | 0.001 |
| $clonal_{HISTORIC}$ | 50 | 0.440 | 0.386 | -- |
| $clonal_{ACCESSION}$ | 8922 | 0.346 | 0.385 | 0.001 |

**Table S7.** The ratio of nonsynonymous to synonymous mutation ($N/S_{mut}$) excluding the subtelomeres. The genomic expectation ($N/S_{exp}$) is calculated from 1000 permutations. For $N/S_{mut}$ values outside of a 95% CI of $N/s_{exp}$, the p-value is the likelihood of a permutation value at least as extreme as the mutation value.

| | | n | $N/S_{mut}$ | $N/S_{exp}$ | p-value |
|---|---|---|---|---|---|
| nonsense | leaf-specific | 1906 | 0.250 | 0.2560 | -- |
| | mosaic | 2725 | 0.094 | 0.216 | 0.005 |
| | $clonal_{HISTORIC}$ | 168 | 0 | 0.051 | -- |
| | $clonal_{ACCESSION}$ | 19085 | 0.117 | 0.211 | 0.001 |
| missense | leaf-specific | 1906 | 6.750 | 3.627 | 0.002 |
| | mosaic | 2725 | 8.094 | 3.428 | 0.002 |
| | $clonal_{HISTORIC}$ | 168 | 1.500 | 3.840 | -- |
| | $clonal_{ACCESSION}$ | 19085 | 1.821 | 2.710 | 0.001 |
| total | leaf-specific | 1906 | 7.000 | 3.912 | 0.002 |
| | mosaic | 2725 | 8.188 | 3.652 | 0.002 |
| | $clonal_{HISTORIC}$ | 168 | 1.500 | 4.318 | -- |
| | $clonal_{ACCESSION}$ | 19085 | 1.938 | 2.924 | 0.001 |

**Table S8.** The ratio of nonsynonymous to synonymous mutation ($N/S_{mut}$) in the subtelomeres. The genomic expectation ($N/S_{exp}$) is calculated from 1000 permutations. For $N/S_{mut}$ values outside of a 95% CI of $N/s_{exp}$, the p-value is the likelihood of a permutation value at least as extreme as the mutation value.

| | | n | $N/S_{mut}$ | $N/S_{exp}$ | p-value |
|---|---|---|---|---|---|
| nonsense | leaf-specific | 2888 | 0.205 | 0.137 | 0.017 |
| | mosaic | 935 | 0.219 | 0.134 | 0.021 |
| | $clonal_{HISTORIC}$ | 50 | 0.000 | 0.134 | -- |
| | $clonal_{ACCESSION}$ | 8922 | 0.157 | 0.159 | -- |
| missense | leaf-specific | 2888 | 2.976 | 2.750 | -- |
| | mosaic | 935 | 2.656 | 2.758 | -- |
| | $clonal_{HISTORIC}$ | 50 | 3.000 | 2.480 | -- |
| | $clonal_{ACCESSION}$ | 8922 | 2.435 | 2.245 | 0.007 |
| total | leaf-specific | 2888 | 3.181 | 2.877 | -- |
| | mosaic | 935 | 2.875 | 2.887 | -- |
| | $clonal_{HISTORIC}$ | 50 | 3.000 | 2.606 | -- |
| | $clonal_{ACCESSION}$ | 8922 | 2.592 | 2.404 | 0.020 |
